# Chromatin remodeling in bovine embryos indicates species-specific regulation of genome activation

**DOI:** 10.1101/2019.12.12.874479

**Authors:** Michelle M Halstead, Xin Ma, Richard M Schultz, Pablo J Ross

## Abstract

The maternal-to-zygotic transition (MZT) is underpinned by wide-spread transcriptomic and epigenomic remodeling that facilitates totipotency acquisition. Factors regulating MZT vary across species and differences in timing of developmental transitions and motif enrichment at accessible chromatin between human and mouse embryos suggest a distinct regulatory circuitry. Profiling accessible chromatin in bovine preimplantation embryos—timing of developmental transitions in bovine closely resembles that in human—indicated that prior to embryonic genome activation (EGA) accessible chromatin is enriched in maternal transcription factor recognition sites, e.g., CTCF, KLFs, NFY, and SP1, echoing observations in humans and mice, and suggesting that a conserved set of maternal factors regulate chromatin remodeling prior to EGA. In contrast, open chromatin established during EGA was primarily enriched for homeobox motifs and showed remarkable similarities between cattle and humans, indicating that cattle could be a more relevant model for human preimplantation development than mice.

## Introduction

Preimplantation development encompasses several critical milestones as embryos progress from fertilization to blastocyst formation. Fusion of the transcriptionally quiescent oocyte and sperm results in a zygote with two haploid pronuclei, which combine during the first round of replication when pronuclear membranes dissolve, allowing maternal and paternal chromosomes to intermingle on the metaphase plate. Subsequent rounds of cleavage ultimately form a blastocyst. However, the cleavage-stage embryo must first complete the maternal-to-zygotic transition (MZT), wherein the embryo assumes control over its own continued development by degrading oocyte-derived products and initiating its own transcriptional program. This dramatic change in gene expression proceeds gradually; minor embryonic genome activation (EGA) results in low levels of transcription in early cleavage-stage embryos^1^, and leads to major EGA, which involves wide-spread activation of embryonic transcription^2^. This shift from maternal dependence to self-sufficiency serves at least three functions: elimination of oocyte-specific messages, replenishment of transcripts that are common to both the oocyte and the embryo, and generation of novel embryonic-specific transcripts.

The MZT also involves wide-spread changes in chromatin structure^3^ and other epigenetic marks, which completely restructures the embryonic epigenome. This chromatin remodeling is necessary to eradicate gamete-specific signatures and establish an open chromatin landscape that supports embryonic transcriptional programs. Specifically, chromatin structure defines the genomic context within which transcriptional machinery can operate, thereby determining the cell-specific gene expression patterns that confer cell identity and function. Following fertilization, the zygotic genome is globally demethylated^4^, and this loss of DNA methylation coincides with global decreases in repressive histone modifications^3,5^. Generally, epigenetic factors linked to relaxed chromatin are more abundant in mouse zygotes, whereas factors implicated in chromatin compaction become more prevalent during EGA^6^, pointing to a highly permissive chromatin landscape in pre-EGA embryos. Indeed, mouse zygotes demonstrate elevated histone mobility^7^, highly dispersed chromatin^8^, and lack chromocenters (congregations of pericentromeric heterochromatin)^9,10^. Moreover, in mouse, 3-D chromatin architecture is largely absent after fertilization, and is then gradually established throughout preimplantation development^11^. This increasing chromatin compaction and organization facilitates long-distance chromatin interactions in later stage embryos^11^. The co-occurrence of genome activation and chromatin remodeling raises an interesting causality dilemma, namely, whether chromatin remodeling is necessary for transcription activation or whether transcription activation leads to chromatin accessibility. Several maternal products prompt transcription initiation by altering chromatin structure^12–14^, and some chromatin compaction occurs in the absence of embryonic transcription^11^. However, inhibiting embryonic transcription pervasively disrupts the establishment of open chromatin during EGA^15,16^, suggesting that EGA and chromatin remodeling are likely interdependent.

In mouse and human preimplantation embryos, accessible sites are gradually established as development progresses^15–20^; however, these open regions demonstrate different motif enrichment patterns, implicating distinct sets of transcription factors (TFs) in either murine (RARG, NR5A2, ESRRB),^15^ or human EGA (OTX2, GSC, POU5F1)^16,18^. Although some TFs appear to regulate EGA in multiple species, i.e. KLFs^15,16,21^, DUX^22–25^, ZSCAN4^26,27^, and CTCF^15,16,28^, it remains unclear if there is any mechanistic conservation across mammals. In fact, the timing of genome activation is highly species-specific: major EGA in mice occurs during the 2-cell stage^1^, in humans^29^ and pigs^30^ during the 4- to-8-cell stage, and in sheep^31^ and cattle^32^ between the 8- and 16-cell stages. The relative timing with which mice activate their genomes could indicate that the mechanism behind murine EGA differs significantly from other species’, which would have significant implications for modeling human preimplantation development. In particular, the timing of bovine EGA more closely resembles that of human EGA, as do global changes to histone PTMs: the active mark trimethylation of lysine 4 on histone 3 decreases in global abundance during human^33^ and bovine EGA^34^, but increases during murine EGA^35,36^. However, the chromatin remodeling events that underscore bovine preimplantation development have yet to be catalogued, and it remains unclear whether the regulation and execution of the MZT in cattle resembles that which has been observed in humans.

To this end, we here describe the chromatin accessibility landscapes of bovine oocytes and preimplantation embryos using the Assay for Transposase Accessible Chromatin (ATAC-seq)^37^. We find that open chromatin is gained progressively throughout development, with promoters enriched for CTCF and NFY motifs gaining accessibility in earlier stages, and distal regions becoming accessible at later stages. Moreover, embryonic transcription was not required for the appearance of promoter open chromatin in 2- and 4-cell embryos, but was absolutely necessary to establish stage-specific and distal open chromatin, especially in 8-cell embryos, indicating that maternal and embryonic products both participate in chromatin remodeling in a complementary fashion. Sequence enrichment in open chromatin revealed that several TFs likely play roles in both bovine and human EGA (OTX2, SP1), whereas regulators specific to murine EGA^15^ demonstrate no enrichment in bovine (RARG, NR5A2, ESRRB), suggesting that cattle may be a more informative model for human preimplantation development. Nevertheless, several TFs (DUX, KLFs, CTCF, NFY) seem to play a role in preimplantation development in all three species, raising the possibility that events leading to EGA may be mechanistically conserved across mammals, whereas the specific transcriptional programs that are activated may differ between species.

## Results and Discussion

### Global dynamics of open chromatin in bovine preimplantation embryos

For each developmental stage, ATAC-seq libraries were prepared from cells derived from three separate oocyte collections. A subset of embryos from each collection was also cultured in the presence of a transcriptional inhibitor (α-amanitin), to interrogate the causal relationship between embryonic transcription and chromatin remodeling (Figure 1a). Between 30 and 87 million non-mitochondrial monoclonal uniquely mapping reads were collectively obtained for each developmental stage, and at least 20 million non-mitochondrial monoclonal reads were collectively obtained for transcription blocked embryos (TBEs) at each stage (Table S1). Genome-wide normalized ATAC-seq coverage demonstrated a striking shift in the open chromatin landscape between 4- and 8-cell embryos (Figure 1b), suggesting that large-scale chromatin remodeling coincided with the main transcriptomic shift observed during major EGA (Figure S1). Similarity between replicates (Table S2) indicated that both the technique and embryo production were robust, generating comparable chromatin accessibility profiles across different rounds of oocyte collection and embryo production (Figure 1c). Reads from replicates were pooled together to obtain greater sequencing depth and maximize power for identifying regions of open chromatin. To gauge changes in chromatin accessibility throughout development, regions of open chromatin, or peaks, were called for each stage of development. To minimize bias from sequencing depth, peaks were called from either 30 million monoclonal uniquely mapped reads when comparing different developmental stages, or 20 million reads when comparing TBE with control embryos.

**Figure 1.**
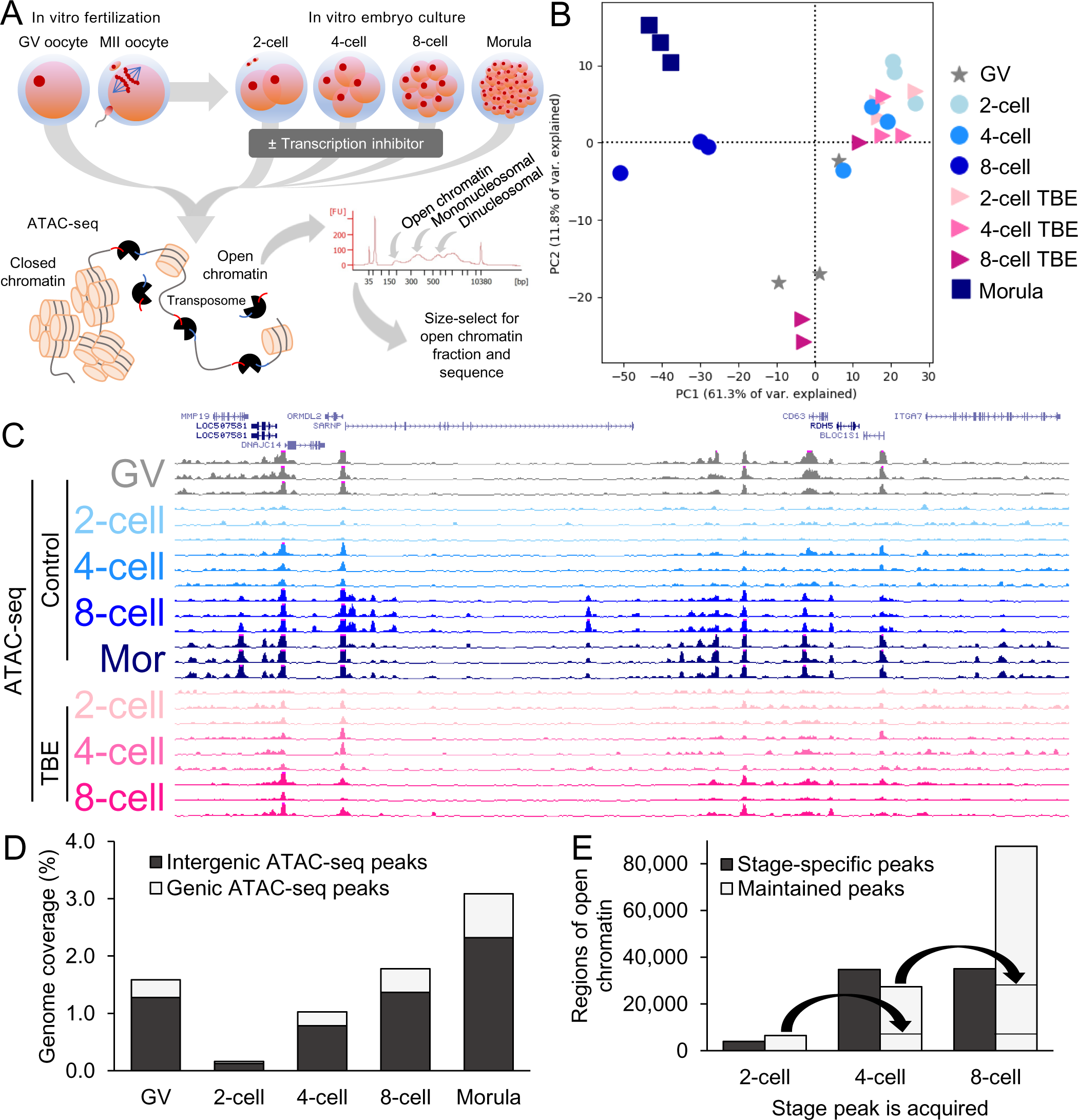
Chromatin accessibility in bovine oocytes and *in vitro* preimplantation embryos. A) Schematic of *in vitro* embryo production and ATAC-seq library preparation. B) Principal components analysis (PCA) of ATAC-seq read depth normalized by reads per kilobase million (RPKM) in 50 bp windows covering the whole genome. C) Normalized coverage (RPKM) of replicate ATAC-seq libraries for each stage of development. D) Proportion of genome covered by genic and intergenic ATAC-seq peaks, called from 30 million reads at each developmental stage. E) Categorization of accessible regions in 2-, 4-, and 8-cell embryos into stage-specific and maintained peaks (accessibility maintained up until the morula stage). Maintained peaks are carried over from latter stages to show cumulative maintained peaks.

As observed in humans and mice^15,16,18–20^, regions of open chromatin were gradually established throughout bovine preimplantation development, with the lowest enrichment for accessible sites in 2-cell embryos (Figure 1d). Rather than reflecting chromatin inaccessibility in 2-cell embryos, this dearth of canonical “open chromatin” probably results from a highly permissive chromatin structure. Indeed, chromocenters are absent from bovine embryos until the early 8-cell stage^9^, indicating a relaxed chromatin configuration in early cleavage-stage embryos. Because assays that employ endonucleases, e.g., ATAC-seq and DNase-seq, depend on increased cutting events at consistently accessible loci, we speculate that genome-wide chromatin relaxation would lead to random cutting events genome-wide, resulting in the observed high background and low enrichment in 2-cell embryos. Attempts to use endonuclease-based methods to profile open chromatin in mouse zygotes^15,20^ and human 2-cell embryos^18,19^ have encountered similar difficulties with low enrichment. As in bovine embryos, chromocenters are also conspicuously absent in pre-EGA mouse embryos^9,10^, and electron microscopy^8^ and fluorescence recovery after photobleaching analysis^7^ also indicate highly dispersed chromatin. Furthermore, a recent study that detected open chromatin based on methylation of accessible GpC sites, rather than endonucleases, found that genome-wide accessibility in human embryos actually decreased from the zygote stage onward^17^. Thus, global chromatin relaxation appears to be a shared characteristic of human, bovine, and murine pre-EGA embryos.

It is tempting to speculate that this “naïve” chromatin state acts as a blank epigenetic slate, which is then gradually compacted and structured to meet the needs of the growing embryo. Indeed, accessible sites were progressively established in 4-cell, 8-cell, and morula stage embryos (Figure 1d). Interestingly, many of these regions were only open transiently at a specific stage, whereas others maintained their accessibility throughout later stages (Figure 1e), suggesting that chromatin remodeling serves two functions: progressive establishment of a “totipotent” chromatin landscape, and transient stage-specific regulation.

### Stepwise remodeling yields stage-specific open chromatin with distinct functionality

To delve further into the potential function of regions that lost or gained accessibility between consecutive stages (Figure 2a), intergenic regions of open chromatin were evaluated for sequence enrichment. Although regulatory regions have not been annotated in cattle, intergenic open chromatin could correspond to enhancers, the activity of which is generally highly tissue-specific. Indeed, distal regions that became accessible at each developmental stage were enriched for different recognition motifs (Table S3). These enriched sequences corresponded to the known binding motifs of several TFs that were either maternally provided or expressed in embryos (Figure S2), demonstrating that the changing chromatin structure subjects each stage of development to distinct regulatory circuitry (Figure 2b).

**Figure 2.**
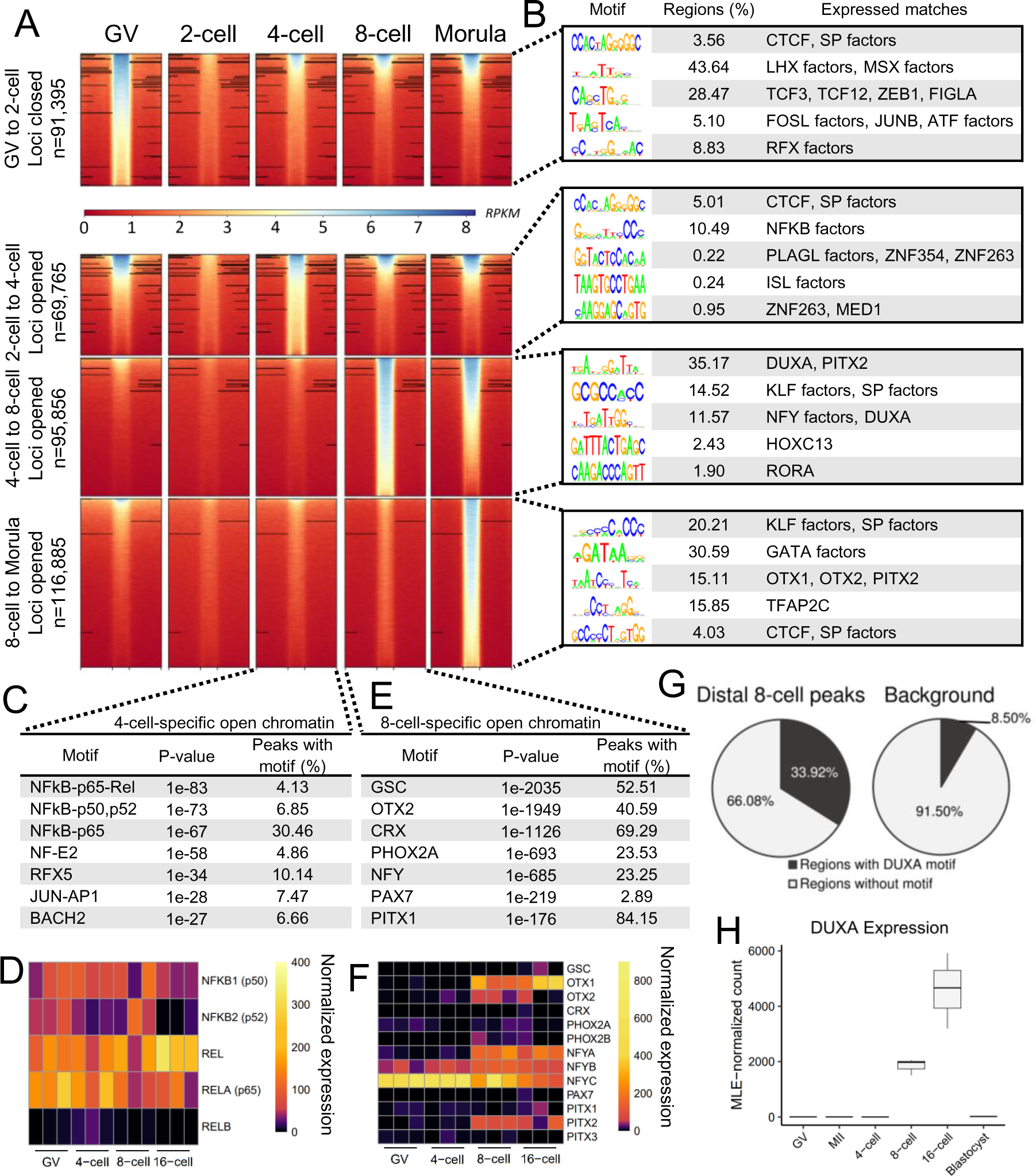
Gradual establishment of open chromatin enriched for regulatory motifs. A) Regions that significantly lost accessibility from the GV oocyte to 2-cell stage, and regions that significantly gained accessibility from the 2- to 4-cell, 4- to 8-cell, and 8-cell to morula stages, according to 30 million reads per stage. Accessibility at each region, scaled to average width ± 1 kb, was determined by normalized ATAC-seq read depth (reads per kilobase million; RPKM), based on 20 million reads per stage. B) Top five enriched *de novo* motifs enriched in intergenic peaks with matching known motifs (match score > 0.6) of TFs that were expressed at the given stage. C) Top seven known motifs enriched in 4-cell specific peaks. D) DESeq2 normalized expression of TFs corresponding to enriched motifs in 4-cell specific open chromatin. E) Top seven known motifs enriched in 8-cell specific peaks. F) DESeq2 normalized expression of TFs corresponding to enriched motifs in 8-cell specific open chromatin. G) Proportion of intergenic 8-cell peaks overlapping the *de novo* motif most closely matching the DUXA motif, relative to background. H) DESeq2 normalized expression of DUXA throughout development. RNA-seq data from Graf *et al* (2014).

The first major transition in chromatin structure mostly involved loss of hyperaccessible sites in 2-cell embryos following fertilization (Figure 2a). Intergenic loci that lost accessibility during this transition (n=54,264) were enriched for the binding motifs of CTCF (an insulator protein implicated in 3D chromatin organization^38^), FIGLA (an oocyte-specific TF), and RFX factors (a highly conserved family of transcriptional repressors^39^; Figure 2b). Interestingly, one quarter of the intergenic regions that closed in 2-cell embryos regained accessibility at the 4-cell stage (n=14,859), and many of these remained open all the way through to the morula stage (n=6,186). Most intergenic regions that closed after fertilization remained inaccessible in embryos, suggesting that they could contribute to oocyte-specific regulation. On the other hand, regions that re-opened in 4-cell embryos seem more likely to participate in house-keeping functions.

The first major gains in accessibility occurred in 4-cell embryos (Figure 2a). About a quarter of the regions that opened during the 4-cell stage remained accessible in both 8-cell embryos and morulae (Figure 1e), although 74% of these had been previously open in GV oocytes (n=15,466). Surprisingly, nearly half of the regions that opened during the 4-cell stage were only transiently accessible (Figure 1e). This 4-cell-specific open chromatin was most significantly enriched for binding motifs of NFkB family members (Figure 2c), which was particularly intriguing because NFkB activation is specifically necessary in 1-cell mouse embryos for development to progress past EGA^40,41^. Although NFkB factors are maternally provided (Figure 2c), they are initially sequestered in the cytoplasm until they translocate into the nucleus at the early 1-cell stage in mice^40,41^ and the 4-to-8 cell stage in cattle^42^. In particular, one of the NFkB subunits capable of activating gene expression – RELA – binds DNA with increased frequency in bovine embryos compared to oocytes^42^, suggesting that NFkB activation of target gene expression could be one of the first events in a cascade leading to major EGA. In fact, one of the few genes with upregulated expression in 4-cell embryos, as compared to MII oocytes, was TRIM8 – a positive regulator of NFkB activity (Figure S3). Additionally, RELA binding sites that were accessible in 4- and 8-cell embryos mark genes that encode key regulators of early preimplantation development, raising the possibility that NFkB binding is involved in their transcription initiation (Figure S4), although the contribution of NFkB signaling to gene expression during minor EGA in cattle has not yet been established.

In contrast to open chromatin in pre-EGA embryos, regions that gained accessibility at the 8-cell stage (Figure 2a) were considerably enriched for the binding motifs of several homeobox TFs, including OTX2, GSC, CRX, PHOX2A, PAX7, and PITX1 (Figure 2b,e), although only OTX1, OTX2, and PITX2 were appreciably expressed during the 8-cell stage (Figure 2f). Most notably, more than a third of the accessible intergenic loci in 8-cell embryos harbored a sequence that most closely matched DUX binding motifs (Figure 2g). The DUX homeoboxes have been extensively implicated in EGA regulation^22–25^, and DUXA is expressed transiently and strongly in bovine embryos during EGA (Figure 2h). The synchronized transcription of DUXA and increased accessibility of its binding sites during bovine development strongly indicates a conserved role for DUX in mammalian preimplantation development, especially considering that DUX family members are highly conserved and specific to placental mammals^43,44^. The factors that regulate DUXA expression in cattle remain unclear. Although maternally provided DPPA2 and DPPA4 induce Dux in mouse embryonic stem cells^45^, only DPPA3 is maternally provided in cattle (Figure S5), and it is unknown if DPPA3 is capable of inducing DUXA expression; however, DPPA3 is maternally provided in mice, cattle, and humans^46,47^, and knockdown of DPPA3 decreases the developmental competency of mouse^48^ and bovine embryos^47^, strongly suggesting that it could activate DUXA expression. Further functional validation will be necessary to determine if DUXA is required for bovine embryogenesis, or if is an important but non-essential regulator of EGA, as in mouse^49–51^.

The expression profile of DUXA in bovine preimplantation development closely mirrored that of another TF implicated in EGA: ZSCAN4 (Figure S6a)^52^. ZSCAN4 is a downstream target of DUX factors in humans^23^ and mice^22,24^, and the coordinated expression of these two factors in cattle certainly suggests a similar mechanism may be at play. Although ZSCAN4 depletion disrupts development past EGA in mice^26^ and cattle^27^, its known binding motifs were not enriched in open chromatin during bovine preimplantation development (Figure S6b-d), suggesting that the binding motif of bovine ZSCAN4 likely differs from those in mice and humans.

Following major EGA, formation of the morula also incurred extensive chromatin remodeling, with even more sites gaining accessibility than during the 8-cell stage (Figure 2a). Compared to earlier stages, intergenic loci that gained accessibility in morulae were primarily enriched for GATA factor binding motifs (Figure 2a; Table S3) – key regulators of trophectoderm establishment and maintenance^53^ – indicating that morulae are already initiating differentiation programs necessary for blastocyst formation.

### Progressive establishment of maintained open chromatin sets the stage for genome activation

Although many regions only experienced stage-specific gains in accessibility, a stable open chromatin landscape was also progressively established after fertilization. As early as the 2-cell stage, regions began to gain accessibility that was maintained until at least the morula stage. These regions of maintained accessibility were heavily enriched for CTCF motifs (Figure 3a; Table S4b), especially those that were first established during the 4-cell stage. CTCF binding delineates chromatin loop boundaries, thus determining the genomic space within which genes can interact with their regulatory elements^38^. Therefore, enrichment of CTCF motifs in maintained peaks could point to a gradual establishment of a stable 3-D chromatin architecture in preparation for major EGA. This proposal is consistent with 3-D chromatin architecture in mouse embryos, which is greatly diminished after fertilization and then gradually re-established throughout preimplantation development, facilitating long-distance chromatin contacts, i.e., promoter-enhancer interactions, in later stage embryos^11^. Indeed, transcription during minor EGA in mouse is primarily driven by proximal promoters, whereas enhancers are dispensable for transcription until major EGA^54^. The global reorganization of 3D chromatin architecture in mouse has also been observed in bovine embryos, wherein gene-rich regions switch from a random distribution to a chromosome-specific distribution during major EGA^55^. Collectively, these results suggest that gradual establishment of 3D chromatin architecture is a conserved feature of pre-EGA embryos, although the mechanisms regulating this restructuring remain unknown.

**Figure 3.**
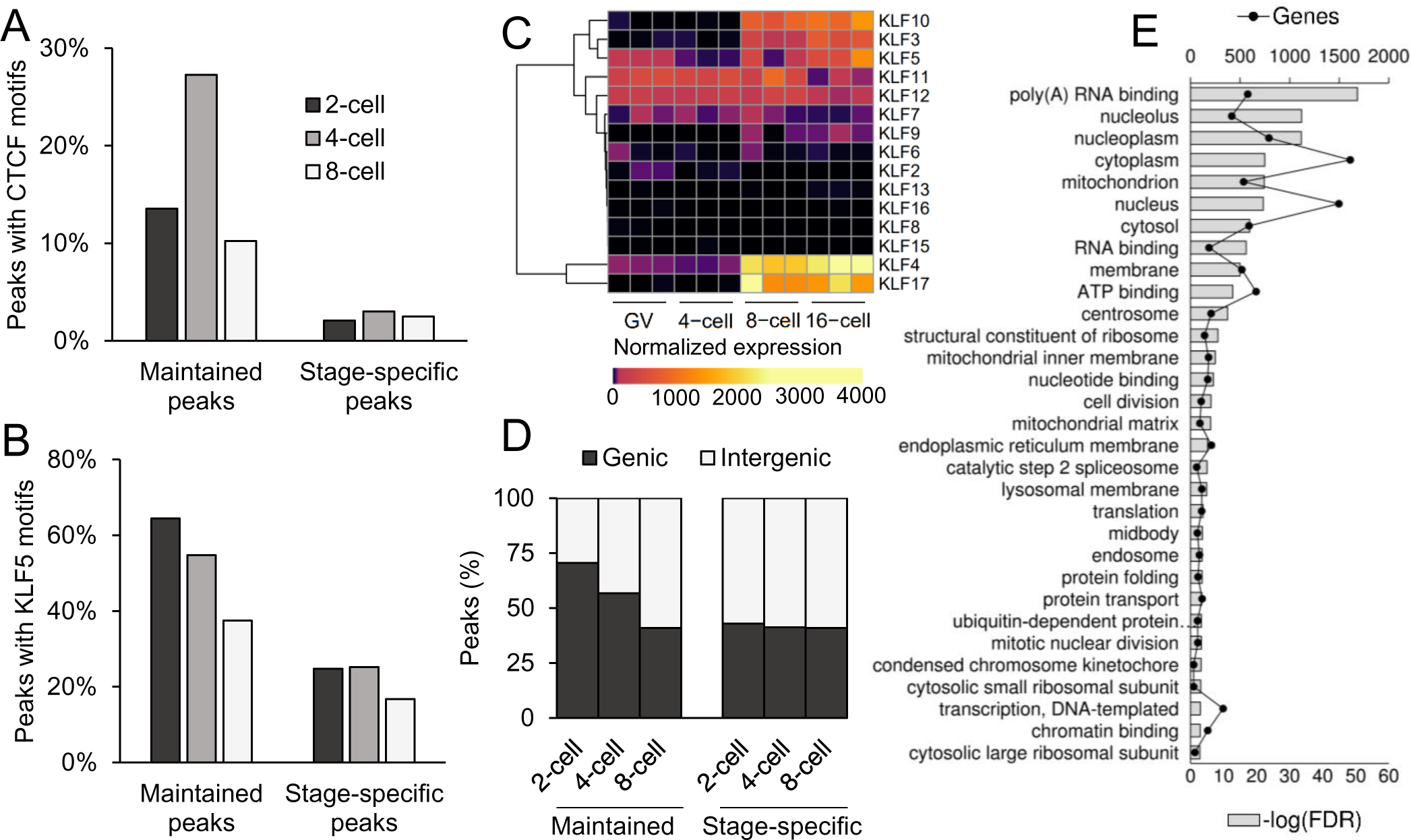
Binding motif enrichment in regions that opened at the 2-, 4-, or 8-cell stages and remained open until at least the morula stage. Proportion of peaks with A) CTCF or B) KLF5 binding motifs. C) DESeq2 normalized expression of KLFs. RNA-seq data from Graf *et al* (2014). D) Proportion of maintained and stage-specific peaks that were genic or intergenic. E) Gene ontology term enrichment of the 9,456 genes that were marked by maintained genic open chromatin that was first established in 2- or 4-cell embryos.

Other than CTCF, maintained peaks were also more enriched for KLF motifs than transiently open sites (Figure 3b). Although most KLFs were not expressed until the 8-cell stage, several KLFs were maternally provided (Figure 3c), including KLF4, a master regulator of induced pluripotent stem cell (iPSC) reprogramming^56^. Recent evidence suggests that KLF4 contributes to reprogramming iPSC by mediating pluripotency-associated enhancer-promoter contacts^57^; thus, the concurrent establishment of open chromatin corresponding to CTCF and KLF binding motifs suggests that pre-EGA embryos are priming a similar mechanism for use during major EGA, especially considering that almost 50% of genes activated during bovine EGA contain KLF motifs in their promoters^21^.

In fact, loci that remained open from the 2- and 4-cell stages onward occurred in genic regions more often than stage-specific open chromatin (Figure 3d) and marked the promoters of genes that were functionally enriched for housekeeping functions (Figure 3e), including 33 of the 51 genes that were upregulated in 4-cell embryos compared to MII oocytes (Figure S3). Moreover, these regions of maintained accessibility were strongly enriched for NFY and SP1 binding sites (Table S4a,b), which is highly reminiscent of chromatin remodeling in mouse embryos, as proximal promoters enriched for NFY are the first regions to gain accessibility^20^. Of note, NFY enhances binding of the pluripotency factors POU5F1, SOX2, and KL4 to their recognition motifs^58,59^, and is clearly involved in murine EGA, as NFY knockdown embryos demonstrated impaired open chromatin establishment and downregulation of gene expression^20^. Similarly, SP1 is essential for early mouse development, with knockout embryos arresting at day 11 of gestation^60^. Intriguingly, human zygotes also demonstrate KLF, SP1, and NFY motif enrichment in open chromatin^19^, overall suggesting that a conserved set of maternal regulators (CTCF, KLFs, SP1, NFY) participates in chromatin remodeling and transcription activation in pre-EGA embryos.

### Both maternal products and embryonic transcription drive chromatin reorganization

Considering that embryonic transcription is extremely limited before the 8-cell stage in cattle, the appearance of open chromatin in pre-EGA embryos suggests that maternal factors participate in chromatin remodeling. To further dissect the maternal contribution to epigenetic reprogramming and EGA, embryos were cultured in the presence of α-amanitin to inhibit POLR2-dependent transcription elongation. Loss of embryonic transcription had a drastic inhibitory effect on the appearance of open chromatin that intensified as development progressed (Figure 4a); 64% of loci that should have opened during the 4-cell stage failed to become accessible without embryonic transcription, and in 8-cell TBE embryos, 96% of loci that should have opened remained closed (Figure 4b), disrupting the chromatin state of key genes, such as KLF4 (Figure 4c), and coinciding with developmental arrest.

**Figure 4.**
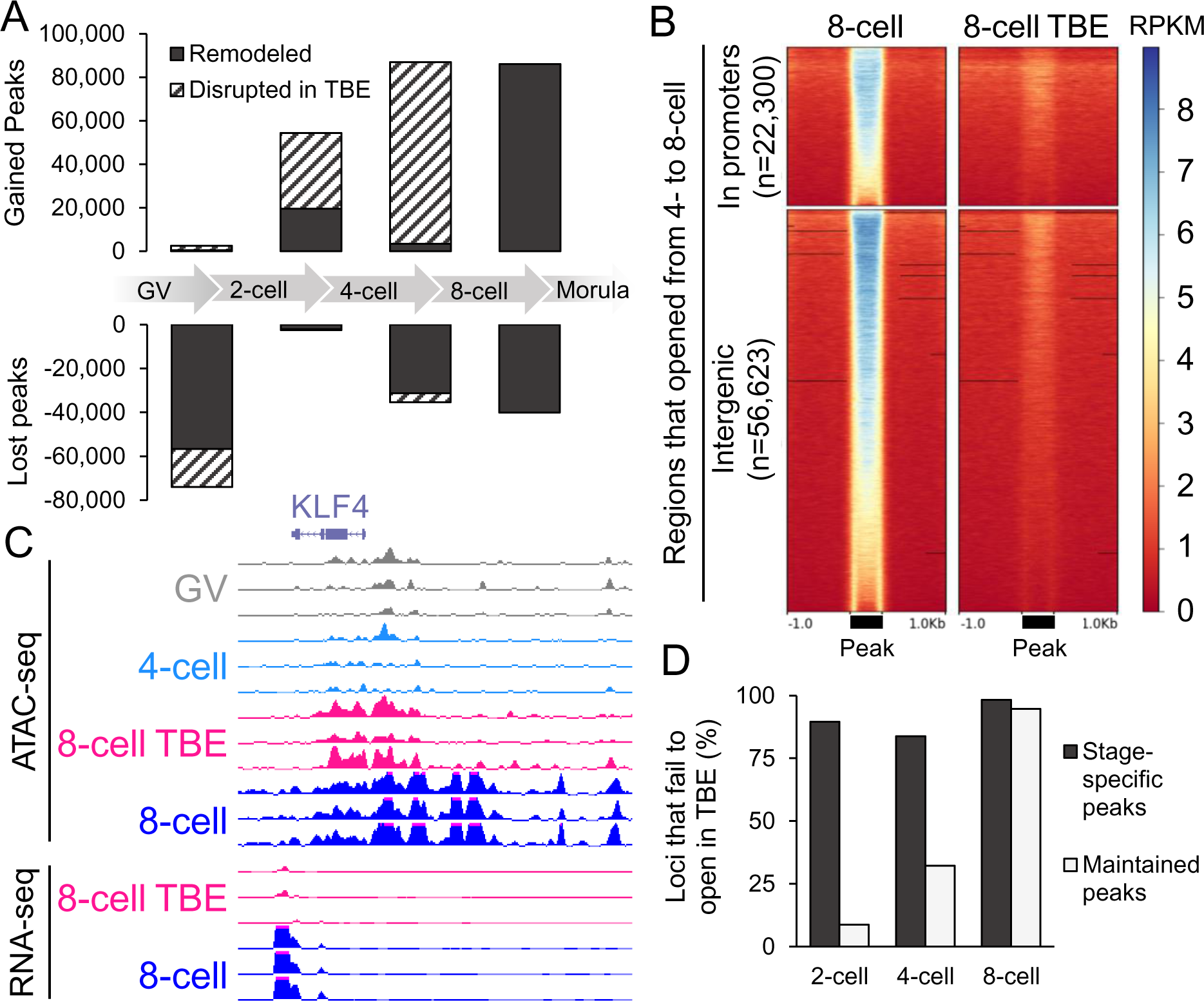
Effect of transcription inhibition on chromatin remodeling. A) Accessibility status of loci that should have opened or closed between consecutive stages in TBEs. ATAC-seq peaks were called based on 20 million reads per stage and categorized as either genic or intergenic. B) Normalized read depth (RPKM) at loci which opened during the 4- to 8-cell transition in 8-cell control and transcriptionally inhibited embryos. Regions scaled to average width ± 1 kb. C) Normalized ATAC-seq and RNA-seq signal (RPKM) in 8-cell and 8-cell TBEs at the KLF4 locus. RNA-seq data from Bogliotti *et al* (2019). D) Proportion of stage-specific and maintained peaks that should appear at each stage, but which fail to open in TBEs.

Interestingly, inhibiting embryonic transcription did not uniformly affect chromatin remodeling genome-wide. Stage-specific open chromatin was preferentially disrupted in TBEs, whereas maintained open chromatin established in 2- and 4-cell embryos appeared even in the absence of embryonic transcription (Figure 4d). Transcription-independent maintained open chromatin marked the promoters of nearly 60% of embryonically-expressed genes (n=1,038/1,784 genes identified from data from Bogliotti *et al* 2019; Figure S7), suggesting that maternal factors actively remodel the local chromatin structure of target genes, possibly priming them for expression later on in development. Nevertheless, the appearance of stage-specific open chromatin almost completely depended on embryonic transcription, indicating that maternal and embryonic factors cooperate in a complementary fashion to establish the appropriate chromatin landscape for activation of embryonic transcriptional programs.

### Open chromatin in preimplantation embryos is enriched for repetitive elements

Evaluation of changes in chromatin structure as they relate to gene expression gives an incomplete perspective of the genome-wide changes that occur during preimplantation development, because the MZT is not just characterized by a shift in the transcriptome but also in the repeatome. Similar to other mammalian species, 44% of the bovine genome is comprised of repeats derived from retrotransposons^61^ – interspersed repeats that are increasingly thought to play major roles in cellular processes and development. Retrotransposons propagate through a ‘copy and paste’ mechanism, and their expression is generally suppressed to avoid deleterious integrations^62^. However, retrotransposons are often actively transcribed in early embryos, and although this phenomenon was recently thought to be nothing more than opportunistic expression by repetitive elements due to an unusually permissive chromatin state in the developing embryo, the activity of some retrotransposons is actually crucial for development^63^. Although the specific mechanisms behind this necessity are still being investigated, transposable elements have been implicated at several regulatory levels, as they can provide binding sites for TFs, allowing them to be co-opted as alternative promoters and enhancers^64^ and participate in 3-D chromatin architecture^65^.

Although repetitive element expression in bovine embryos was reported a decade ago using a cDNA array^66^, a complete catalogue of repeat transcription throughout bovine preimplantation development was lacking. To address this gap in knowledge, available RNA-seq data^67^ were assessed for expression of repetitive elements. Importantly, these libraries were not subjected to polyA selection. As has been observed in the mouse and human^68,69^, expression and accessibility of repetitive elements throughout bovine preimplantation development was highly stage-specific and dynamic (Figure 5a; Figure S8).

**Figure 5.**
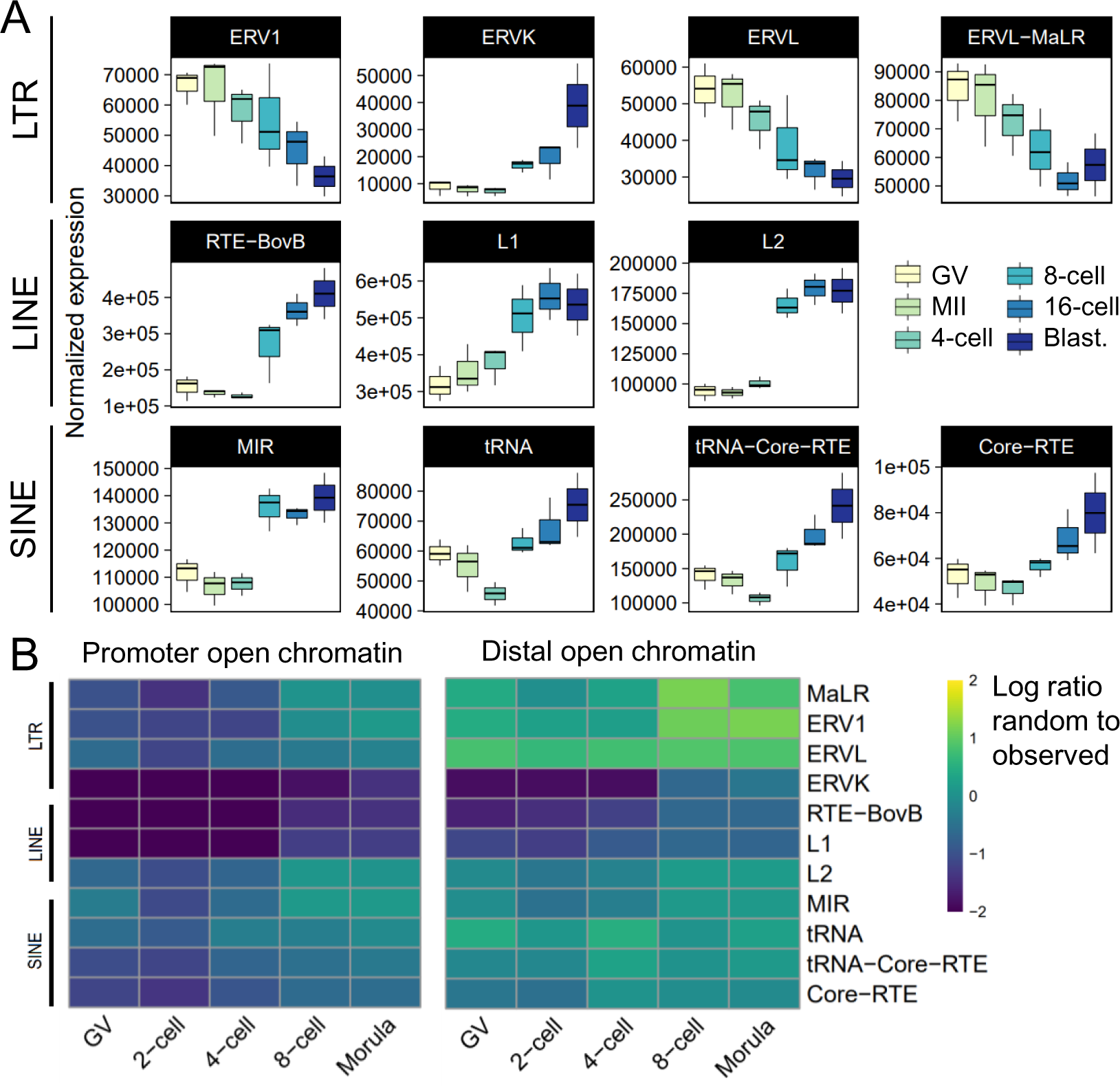
Expression and accessibility dynamics of repeat families during bovine preimplantation development. A) DESeq2 normalized expression profiles of select LTR, LINE, and SINE repeat families. RNA-seq data from Graf *et al* (2014). B) Enrichment of several transposable element families in ATAC-seq peaks, called from 30 million reads. Promoter peaks fell within the 2 kb region upstream of transcription start sites (TSS). Intergenic peaks did not overlap the 2 kb regions upstream of TSS, exons, or introns.

Non-long terminal repeat (non-LTR) retrotransposons – long interspersed elements (LINEs) and short interspersed elements (SINEs) – are increasingly transcribed during human preimplantation development^17,68^. In cattle, LINEs also demonstrated increasing expression (Figure 5a) and accessibility (Figure 5b) starting during the 8-cell stage. Although the function of LINE elements in bovine preimplantation embryos has yet to be established, their activation is crucial for mouse development: perturbing LINE expression in mouse preimplantation embryos causes developmental arrest at the 2-cell stage and perturbs gene expression^70^. Moreover, LINE-1 activation may regulate global chromatin accessibility in mouse embryos^71^. Of the SINE families, mammalian-wide interspersed repeats (MIR) expression and accessibility patterns most echoed those of L2 LINE elements, with increased expression and accessibility starting at the 8-cell stage, suggesting that these elements could be acting as enhancers or promoters.

In particular, LTR activation is a key feature in human^68^, mouse^69^, and bovine preimplantation development^66^. Of these, mammalian LTRs and endogenous retroviral elements (ERVL) were increasingly enriched in open chromatin starting during the 8-cell stage (Figure 5b), although their transcript abundance dropped throughout development (Figure 5a), indicating that other mechanisms likely regulate repeat expression, e.g., DNA methylation or histone modifications^72,73^. Only endogenous retroviral K elements (ERVK) demonstrated both increasing expression (Figure 5a) and moderately increased prevalence in distal open chromatin during the 8-cell stage (Figure 5b), suggesting that ERVK elements function as regulatory elements, as observed in human preimplantation embryos^74^.

Mounting evidence suggests that specific types of LTR retrotransposons, especially intact elements, play pivotal roles in early development. During bovine preimplantation development, the most expressed retrotransposons were ERV1-1_BT and ERV1-2_BT (Figure 6a), as has been observed previously^66^. Upon further inspection, intact ERV1-1_BT elements often co-occurred with MER41_BT repeats in a specific sequence, which demonstrated a highly reproducible pattern of transcription at ERV1-1_BT elements and chromatin accessibility at MER41_BT elements (Figure S9a). Furthermore, MER41_BT elements that were accessible in 8-cell embryos were enriched for the binding motifs of several pluripotency factors, including POU5F1, NFY, KLF4, OTX2, and TEAD (Figure S9b), suggesting that pluripotency factors are driving transposon expression.

**Figure 6.**
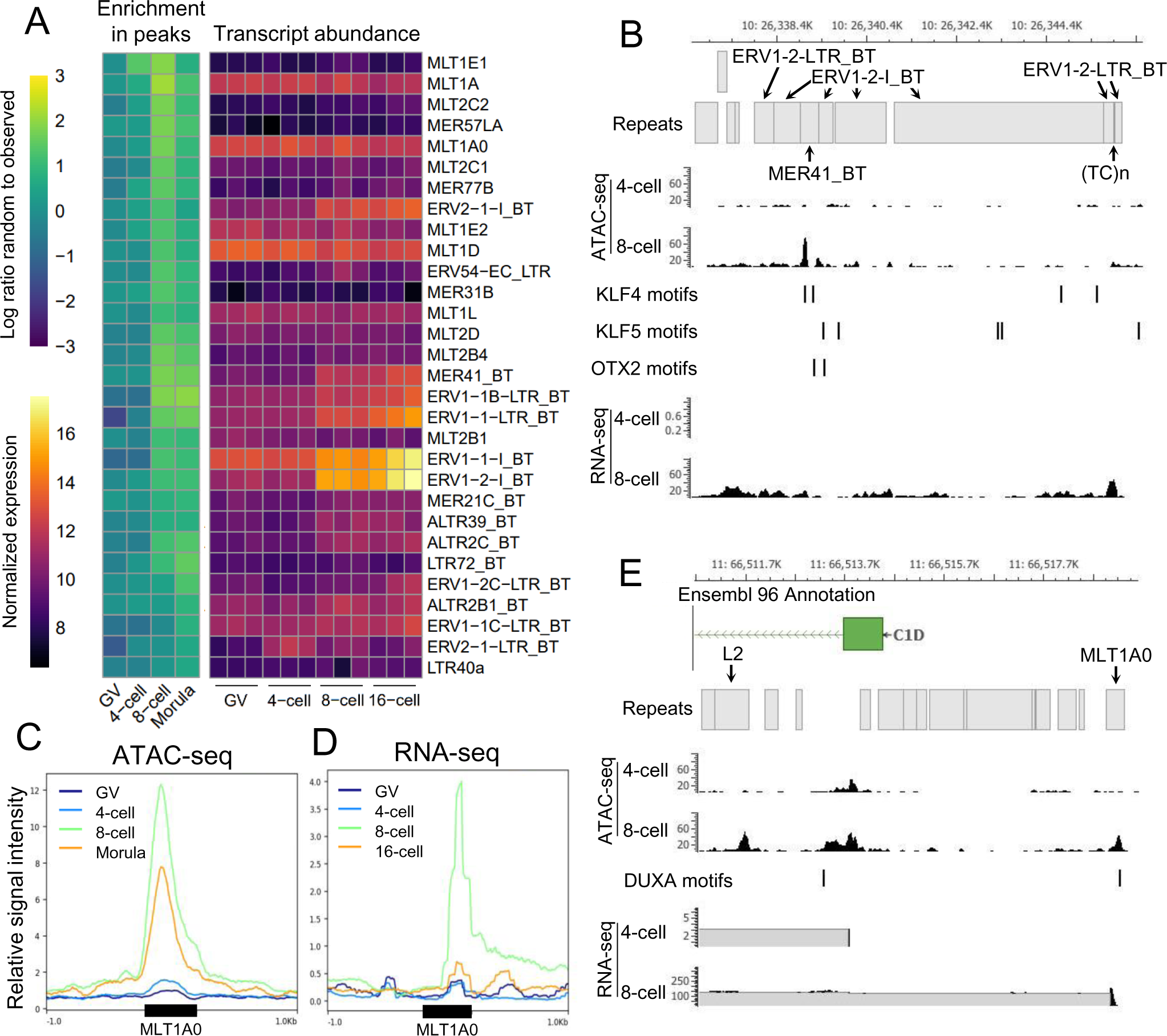
Activity of LTR elements in bovine preimplantation embryos. A) Enrichment of LTR elements in open chromatin in 4-cell, 8-cell, and morula-stage embryos, and expression of the same LTR elements. Variance-stabilized normalized expression shown for three replicates per stage. B) ATAC-seq and RNA-seq read coverage at a highly expressed ERV1-2_BT locus. ATAC-seq tracks show coverage from 30 million reads per library; RNA-seq tracks show combined coverage from three replicates. TF motifs predicted from JASPAR binding motifs MA0039.3 (KLF4) and MA0712.1 (OTX2), respectively. C) Average normalized ATAC-seq and D) RNA-seq signal (RPKM) at MLT1A0 repeats overlapped by 8-cell intergenic open chromatin harboring DUXA motifs, predicted from JASPAR motif MA0468.1. E) Co-option of an accessible MLT1A0 element with a DUXA binding motif as an alternative promoter upstream of the C1D locus. RNA-seq data from Graf *et al* (2014).

DUX has also been implicated in driving the expression of intact ERVL elements in human and mouse pre-EGA embryos^19,22–24^. Specifically, in human embryos DUX4 appears to bind MLT2A1 elements – primate-specific LTRs that flank human ERVL –activating their expression^19^. Considering that intergenic open chromatin was especially enriched for DUX binding sites in 8-cell bovine embryos, it seemed likely that these sites would also correspond to retrotransposons. Indeed, several MLT elements were enriched in 8-cell open chromatin harboring DUX motifs (Table S5), suggesting that bovine DUX may bind and regulate LTRs with sequence similarity to primate-specific MLT2A1 elements. Nearly all LTRs that were enriched in 8-cell open chromatin with DUX binding sites demonstrated dynamic expression profiles throughout development (Figure S10). In particular, increasing accessibility at MLT1A0 elements harboring DUX recognition sites (Figure 6c) was mirrored by a sharp increase in transcription in 8-cell embryos (Figure 6d).

Whether the transcripts derived from LTRs are required for bovine development or are simply a result of opportunistic expression remains to be established. However, evidence in other species suggests that retrotransposon-derived regulatory elements are often co-opted by the embryo as promoters and enhancers^64^: a phenomenon which appears to extend to bovine embryos, as ATAC-seq and RNA-seq suggest that MLT1A0 elements are co-opted as alternative promoters at several loci, including CD1 (Figure 6e), ZNF41, and LPIN2. As such, an interesting balance appears to exist between repetitive elements and the embryo, wherein the repeatome leverages the embryo’s existing regulatory network to drive transposition, while simultaneously providing new regulatory elements and TF binding sites that the embryo co-opts to drive the expression of its own transcriptional program.

### A model for mammalian genome activation

Several lines of evidence suggest that the regulatory circuitry responsible for the MZT may differ between mammals. First, EGA occurs in a highly species-specific fashion, with the major wave of transcription in mice during the 2-cell stage^1^, in humans during the 4-to-8-cell stage^29^, and cattle during the 8-to-16-cell stage^32^. Second, the expression patterns of repetitive elements are not only highly-stage specific, but also species-specific; primate-specific and murine ERVL elements are strongly expressed during human and mouse preimplantation development, respectively^19,22–24^, whereas ERV1 elements were most prominently activated in bovine embryos. Finally, the maternal programs in human and mice are divergent; human maternal programs conspicuously lack the murine maternal effect transcripts POU5F1, HSF1, and DICER1, and are functionally enriched for translational processing^46^, reflecting the need to translate maternal messages during the extended period between fertilization and human EGA.

The relaxed chromatin structure in early preimplantation embryos provides a unique regulatory context for maternal factors, which are essential to support cleavage stage-embryos prior to genome activation, especially in species where EGA is delayed for several cell divisions, e.g., humans and cattle. Whereas the condensed chromatin structure in somatic cells generally restricts DNA-binding proteins to regions of open chromatin, the dispersed chromatin in 2-cell bovine embryos may allow maternal factors to opportunistically and pervasively bind their recognition motifs. Indeed, several maternal factors appear to participate in chromatin restructuring in pre-EGA embryos, leading to open chromatin establishment at promoters enriched for NFY and SP1 binding sites, as well as CTCF and KLF motifs. Although not maternally provided, the DUX family also appears to play a conserved role in genome activation. In humans and mice, DUX factors have been implicated in chromatin remodeling^23,25^ and transcription activation of cleavage-stage genes^23^. We find a similar pattern of DUX expression in bovine embryos, as well as increased accessibility of DUX binding sites around the 8-cell stage, suggesting that DUX may also modulate gene expression and chromatin accessibility during bovine EGA.

Although the chromatin landscape changes markedly upon major EGA in bovine, human^16,17,19^, and mouse^15,20^ embryos, the regulatory circuitry that is active during this stage in development appears to differ significantly between humans and mice^16^, which suggests that regulation of mammalian EGA is highly species-specific. To identify and compare putative regulators of mammalian EGA, intergenic regions that were accessible during major EGA in bovine, human, and mouse embryos were evaluated for binding motif enrichment of actively expressed TFs. Comparing EGA regulatory circuitry between species reflects a stark divergence in regulatory and transcriptional programs that clearly separates humans and cattle from the mouse (Figure S11). Compared to mouse 2-cell embryos, open chromatin in bovine and human 8-cell embryos demonstrated remarkably similar patterns of sequence enrichment (Figure 7). In cattle and humans, SP1, OTX2, and NFY were particularly implicated during major EGA, whereas NR5A2, RARG, and ESRRB were solely enriched in mouse embryos.

**Figure 7.**
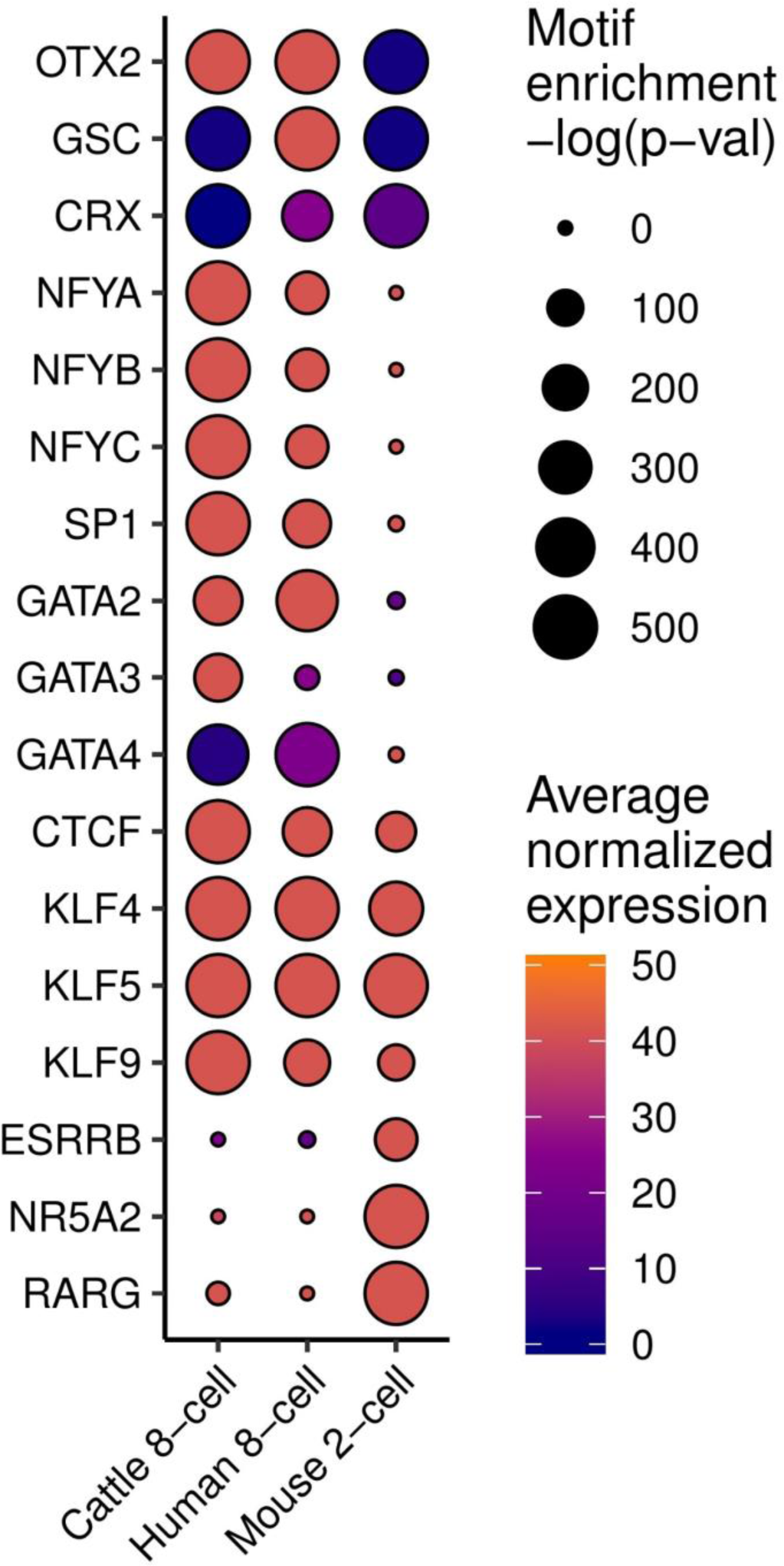
Inference of key regulatory factors during EGA in cattle, human, and mouse, based on enrichment of TF binding motifs in open chromatin and expression of the corresponding TFs. Bovine RNA-seq data from Graf *et al* (2014); human ATAC-seq and RNA-seq data from Wu *et al* (2018); mouse ATAC-seq and RNA-seq from Wu *et al* (2016).

Nevertheless, it is unclear if the regulatory factors that are enriched in open chromatin during major EGA are regulators of EGA or simply products of it. For instance, although NR5A2 is enriched in 2-cell mouse embryos, it is an early regulator of inner cell mass and trophectoderm programs and is not essential for genome activation^15^. Similarly, OTX2 is essential for neuronal lineage specification in mice^75^, and has been implicated in the transition from naïve to primed pluripotency^76^. However, OTX2 protein is clearly present in human and marmoset zygotes^46^, suggesting that this homeobox TF may play an as-of-yet undetermined role in EGA. Although several TFs appeared to only be important in mouse, or in cattle and humans, KLFs were substantially enriched during EGA in all three species. Considering the well-established role of KLFs in somatic cell reprogramming and establishment of pluripotency in multiple species^56,77^, KLFs may play a conserved role in the MZT. Although future research will be necessary to elucidate the function of specific regulatory factors, the high consistency between cattle and humans, both with respect to the timing of EGA and the regulatory circuitry that accompanies it, strongly suggests that cattle are a more appropriate model system for human preimplantation development than mouse.

### Conclusions

Sweeping changes to chromatin structure during bovine preimplantation development suggest that 2-cell embryos are characterized by globally decondensed chromatin, which is gradually compacted as development progresses, echoing similar observations in humans and mice. In particular, it is tempting to speculate that a conserved set of maternal factors establish basal promoter accessibility and the necessary chromatin architecture for enhancer-promoter interactions that will drive gene expression during major EGA (Figure 8). However, the open chromatin landscape during major EGA clearly distinguished mice from cattle and humans, suggesting that whereas maternal regulation of EGA may be conserved across mammals, the transcriptional programs that are subsequently activated have diverged substantially. Practically, this difference suggests that human development may be better modeled in cattle than in mice. Nevertheless, the factors appear to regulate the MZT in cattle, humans, and mice certainly warrant further investigation and validation, which will provide invaluable insight into the regulatory framework that governs successful preimplantation development.

**Figure 8.**
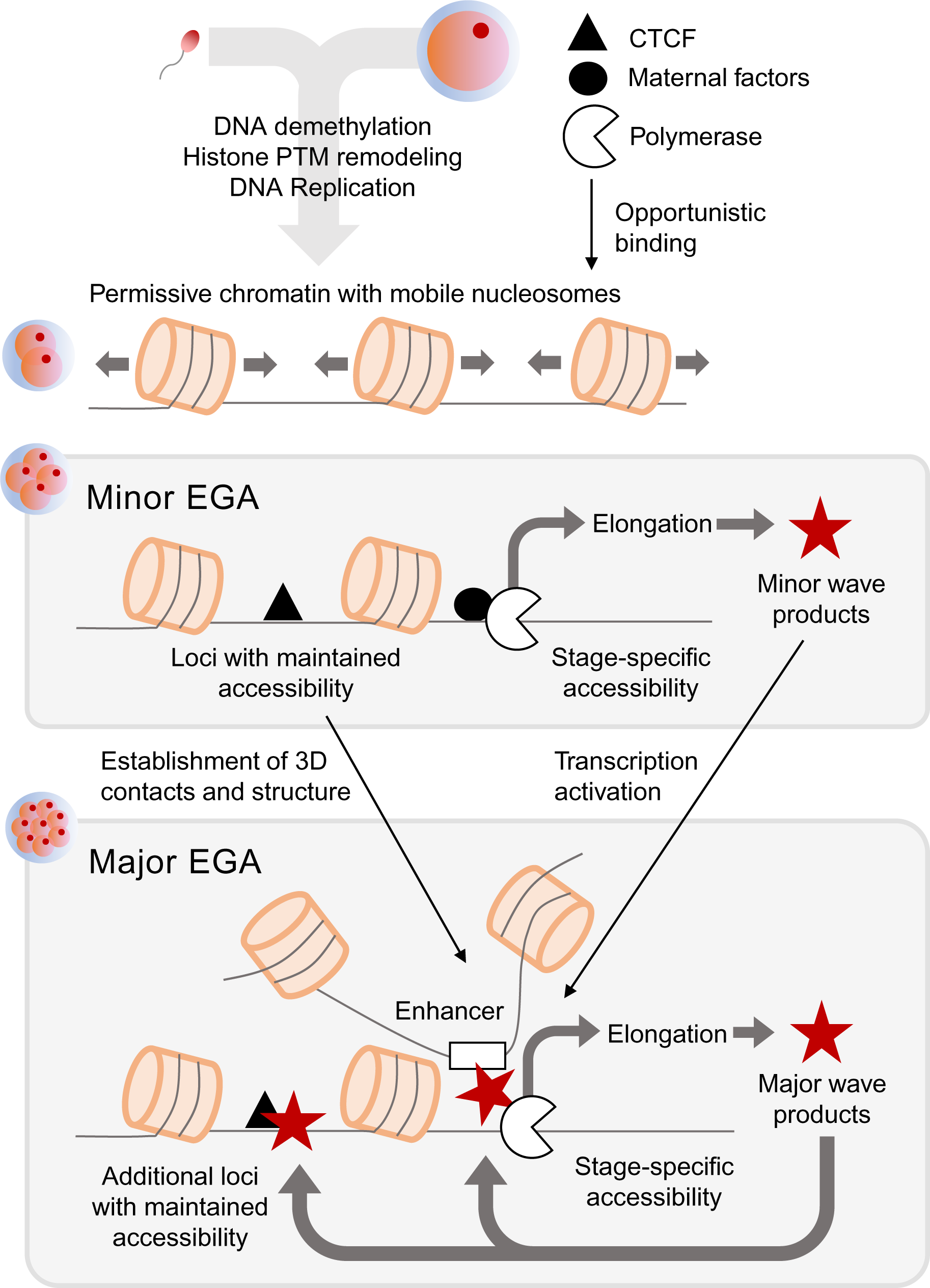
Potential mechanistic model depicting events leading to major EGA. We postulate that chromatin structure is globally decondensed following fertilization, allowing opportunistic binding of maternal factors which initiate a minor wave of transcription and begin to establish 3D chromatin architecture. This sets the stage for major EGA, wherein maternal products, minor EGA products, and promoter-enhancer contacts collectively regulate the first major wave of gene expression and continue to refine 3D chromatin structure.

## Materials and Methods

### Oocyte collection and maturation

Ovaries were procured from a local abattoir and transported to the laboratory in a warm saline solution. Follicles measuring 2-10 mm were aspirated to obtain cumulus oocyte complexes (COCs). Only COCs with healthy layers of cumulus cells were selected for maturation. These were washed in collection medium (6:4 M199 (Sigma M7653)): SOF-Hepes, supplemented with 2% fetal bovine serum (FBS; Hyclone/Thermo Scientific) and transferred to maturation medium (modified M199 medium (Sigma M2154)) supplemented with ALA-glutamine (0.1 mM), sodium pyruvate (0.2 mM), gentamicin (5 μg/ml), EGF (50 ng/ml), oFSH (50 ng/ml), bLH (3 μg/ml), cysteamine (0.1 mM), and 10% FBS.

### In vitro fertilization and embryo culture

After COCs matured for 24 h, MII oocytes were washed in SOF-IVF medium (107.7 mM NaCl, 7.16 mM KCl, 1.19 mM KH^2^PO^4^, 0.49 mM, MgCl^2^, 1.17 mM CaCl^2^, 5.3 mM sodium lactate, 25.07 mM NaHCO^3^, 0.20 mM sodium pyruvate, 0.5 mM fructose, 1X non-essential amino acids, 5 μg/ml gentamicin, 10 μg/ml heparin, 6 mg/ml fatty acid-free (FFA) BSA) and transferred to drops of SOF-IVF medium under mineral oil. Frozen semen from a Holstein bull was thawed, and 10^6^ sperm/ml were added to drops with MII oocytes, which were incubated at 38.5°C for 12-18 h. Zygotes were then removed from fertilization medium, and cumulus cells were removed by vortexing for 5 min in SOF-Hepes medium. Zygotes were then transferred to culture medium (KSOMaa supplemented with 4mg/mL BSA) under mineral oil, and incubated at 38.5°C in 5% CO^2^, 5% O^2^, and 90% N^2^. If embryos were to be transcriptionally inhibited, the culture medium was supplemented with α-amanitin (50 μg/ml) on day one. On day three, the culture medium was supplemented with 5% stem-cell qualified FBS (Gemini Bio 100-525). Blastocyst development was evaluated at 7 days post-insemination (dpi).

### Collection of oocytes and preimplantation embryos for ATAC-seq

Oocytes and embryos were collected for ATAC-seq library preparation from three separate collections per developmental stage. Embryos intended for collection at the 2-cell, 4-cell, or 8-cell stages were divided into two groups, one of which was supplemented with α-amanitin, and cultured simultaneously. Germinal vesicle-stage oocytes were collected for ATAC-seq prior to maturation. Preimplantation embryos were collected at the 2-cell (30-32 h post-insemination), 4-cell (2 dpi), 8-cell (3 dpi), and morula stages (5 dpi).

### ATAC-seq library preparation

Oocytes or embryos (a minimum of 500 cells) were treated with pronase (10 mg/ml) to completely remove the *zona pellucida* and washed with SOF-Hepes on a warming plate. Cells were then transferred to 1 ml cold SOF-Hepes, and centrifuged at 500 rcf, 4°C, for 5 min. Morulae were subjected to additional vortexing for 3 min in cold ATAC-seq lysis buffer (10 mM Tris-Cl, pH 7.4, 10 mM NaCl, 3 mM MgCl^2^ and 0.1% IGEPAL CA-630). Cell pellets were then resuspended in 1 ml cold ATAC-seq cell lysis buffer and centrifuged at 500 rcf, 4°C, for 10 min. Nuclear pellets were then resuspended in 50 μl transposition reaction mix (25 μl TD buffer (Nextera DNA Library Prep Kit, Illumina), 2.5 μl TDE1 enzyme (Nextera DNA Library Prep Kit, Illumina), 22.5 μl nuclease-free H^2^O) and incubated for 60 min at 37°C, shaking at 300 rpm. The transposase, which is loaded with Illumina sequencing adapters, cuts DNA where it is not sterically hindered and simultaneously ligates adapters, effectively producing a library in one incubation step. Transposed DNA was purified with the MinElute PCR purification kit (Qiagen, Hilden, Germany) and eluted in 10 μl buffer EB. Libraries were then PCR amplified: 50 μl reactions (25 μl SsoFast Evagreen supermix with low ROX (Bio-Rad, Hercules, CA), 0.6 μl 25 μM custom Nextera PCR primer 1, 0.6 μl 25 μM custom Nextera PCR primer 2 (for a list of primers, see Buenrostro *et al* (2015)^78^), 13 μl nuclease-free H^2^O, and 10 μl eluted DNA) were cycled as follows: 72°C for 5 min, 98°C for 30 s, and then thermocycling at 98°C for 10 s, 63°C for 30 s and 72°C for 1 min. Libraries from GV oocytes, 2-cell, and 4-cell embryos were thermocycled for 13 cycles, and 8-cell and morulae libraries were thermocycled for 11 cycles. PCR-amplified libraries were again purified with the MinElute PCR purification kit and eluted in 10 μl buffer EB. Libraries were then evaluated for DNA concentration and nucleosomal laddering patterns using the Bioanalyzer 2100 DNA High Sensitivity chip (Agilent, Palo Alto, CA). Expected nucleosomal laddering was evidenced by the presence of both small fragments, corresponding to hyper-accessible DNA that was frequently transposed, and larger fragments, corresponding to DNA that was wrapped around one or more nucleosomes. This study focused on mapping open chromatin; therefore, the sub-nucleosomal length fraction of each library (150-250 bp) was size selected using the PippinHT system (Sage Science, Beverly, MA) with 3% agarose cassettes. Size-selected libraries were run on a Bioanalyzer DNA High Sensitivity chip to confirm size-selection and determine DNA concentration. Final libraries were then pooled for sequencing on the Illumina NextSeq platform to generate 40 bp paired end reads.

### ATAC-seq read alignment and peak calling

Raw sequencing reads were trimmed with Trim_Galore, a wrapper around Cutadapt (v0.4.0)^79^, to remove residual Illumina adapter sequences and low quality (q<20) ends, keeping unpaired reads and reads 10 bp or longer after trimming. Trimmed reads were then aligned to either the GRCm38 (mouse), GRCh38 (human), or ARS-UCD1.2 (cattle) assemblies using BWA aln (-q 15) and sampe^80^. PCR duplicates were removed with PicardTools (v2.8.1), and mitochondrial and low-quality alignments (q<15) were removed with SAMtools (v1.7)^81^. Alignments from biological replicates from each stage were merged and randomly subsampled to equivalent depth with SAMtools for detection of open chromatin. To determine which regions of the genome demonstrated significant enrichment of ATAC-seq signal, broad peaks were called with MACS2 (v2.1.1)^82^, using a q-value cutoff of 0.05, and settings *--nomodel --shift -100 --extsize 200*.

### RNA-seq alignment and gene expression quantification

Raw sequencing reads were trimmed with Trimmomatic (v0.33)^83^. Low-quality leading and trailing bases (3 bases) were clipped, and Illumina adapter sequences were removed, allowing 2 seed mismatches, a palindrome clip threshold of 30, and a simple clip threshold of 10. Sliding window trimming was conducted with a window size of 4 bases, and a quality threshold of 15. Reads 36 bases or longer were retained after trimming. Trimmed reads were aligned to either the GRCm38 (mouse), GRCh38 (human), or ARS-UCD1.2 (cattle) assemblies with STAR (v2.7.2a)^84^ with options *–outFilterScoreMinOverLread 0.85* and *–seedSearchStartLmax 30*. Low quality alignments (q<5) were removed with SAMtools. Raw counts were calculated for genes in the Ensembl 96 annotations for each species with summarizeOverlaps, from the R package GenomicAlignments (v1.18.1)^85^, using ‘Union’ mode and allowing fragments for paired end data. Gene counts were MLE-normalized using the DESeq2 R package (v1.22.2)^86^ and submitted to the variance stabilizing transformation for some analyses. DESeq2 was also used for differential expression analysis, with genes demonstrating a logFC > 2 and an adjusted p-value < 0.05 considered differentially expressed.

### Repeat expression quantification

Trimmed RNA-seq reads were aligned to the ARS-UCD1.2 genome assembly with STAR with options *– outFilterMultimapNmax 100, –winAnchorMultimapNmax 100,* and *–twopassMode Basic*. Raw expression values for individual repetitive elements were calculated for repeats in the RepeatMasker annotation for the ARS-UCD1.2 assembly (downloaded from the UCSC Genome Browser) using TEtoolkit (v2.0.3)^87^ in ‘multi’ mode, which improves quantification of transposable elements transcripts by including ambiguously mapped reads. Raw expression values were MLE-normalized using DESeq2.

### Comparison of replicate libraries and ATAC-seq and RNA-seq signal at regions of interest

For both ATAC-seq and RNA-seq data, alignments were converted to bigwig format using bamCoverage from the DeepTools suite^88^, which binned the genome into 50 bp windows and calculated normalized signal (reads per kilobase million; RPKM) in each window. The plotPCA function from DeepTools was then used to generate principal components plots, with options *–transpose* and *–log2*. The plotCorrelation function from DeepTools was used to calculate the Spearman correlation coefficient between replicate libraries, based on genome-wide normalized coverage. To assess average accessibility or expression at genomic intervals of interest, average ATAC-seq or RNA-seq normalized signal from bigwig files was visualized using the Deeptools plotHeatmap function.

### Comparison and classification of ATAC-seq peaks

Peak sets from different stages were compared using the BEDtools intersect function^89^, requiring a minimum of 1 bp overlap to consider a peak shared by both sets. Similarly, peaks were classified as genic if they overlapped either the 2 kb region upstream of a transcription start site (TSS), exons, or introns by 1 bp. Otherwise, peaks were considered intergenic.

### Motif enrichment

Genomic regions were evaluated for binding motif enrichment using the findMotifsGenome.pl script from HOMER (v4.8)^90^, using the exact sizes of the input genomic intervals (*–size given*). The most significant known or *de novo* motifs were reported, based on p-value. Known motifs that matched significantly enriched *de novo* motifs were reported if their match score exceeded 0.6.

### Genome-wide motif location prediction

Position-weight matrices were downloaded from the JASPAR database for TFs of interest^91^. Using the FIMO tool from the MEME suite (v5.0.4)^92^, TF motif locations (p < 1e-4) were predicted genome-wide in the ARS-UCD1.2 genome assembly.

### Repeat class enrichment in genomic intervals

To determine if repetitive elements (either individual elements, families, or classes) were enriched in open chromatin, the number of ATAC-seq peaks overlapping a set of repetitive elements was compared to randomized intervals (ATAC-seq peak locations shuffled with BEDtools shuffle function) overlapping the same set of repetitive elements, yielding a log ratio of random to observed.

### Functional annotation enrichment analysis

Gene sets were submitted to DAVID (v6.8)^93,94^ to identify enriched biological functions. Gene ontology terms with a false discovery rate (FDR) < 0.05 were reported.

## Acknowledgements

This work was supported by a grant from NIH-NICHD (R01HD070044) to P.J.R. and R.M.S. Semen used for *in vitro* embryo production was provided by Semex (Madison, WI). M.M.H. was supported by USDA NIFA National Need Fellowship #20143842021796 and an Austin Eugene Lyons Fellowship. The authors thank the entire Ross laboratory for contributions to embryo production and lively discussion.

## Author contributions

M.M.H., R.M.S. and P.J.R conceived and designed the experiments; X.M. and M.M.H performed the experiments; M.M.H., R.M.S. and P.J.R wrote the manuscript.

## Data availability

The following published data sets were used, and accessed through the NCBI GEO repository: for bovine oocytes and *in vitro* produced embryos, raw RNA-seq data were downloaded from accession number GSE52415^67^, and mouse and human preimplantation embryo ATAC-seq and RNA-seq data, raw sequencing files were downloaded from accession numbers GSE66390^15^ and GSE101571^16^, respectively. The ATAC-seq data produced in this study is available via the NCBI SRA repository under the SRA accession number PRJNA595394.

