## Supplementary files for "Chromatin remodeling in bovine embryos indicates species-specific regulation of genome activation"

Supplementary Figures and Tables

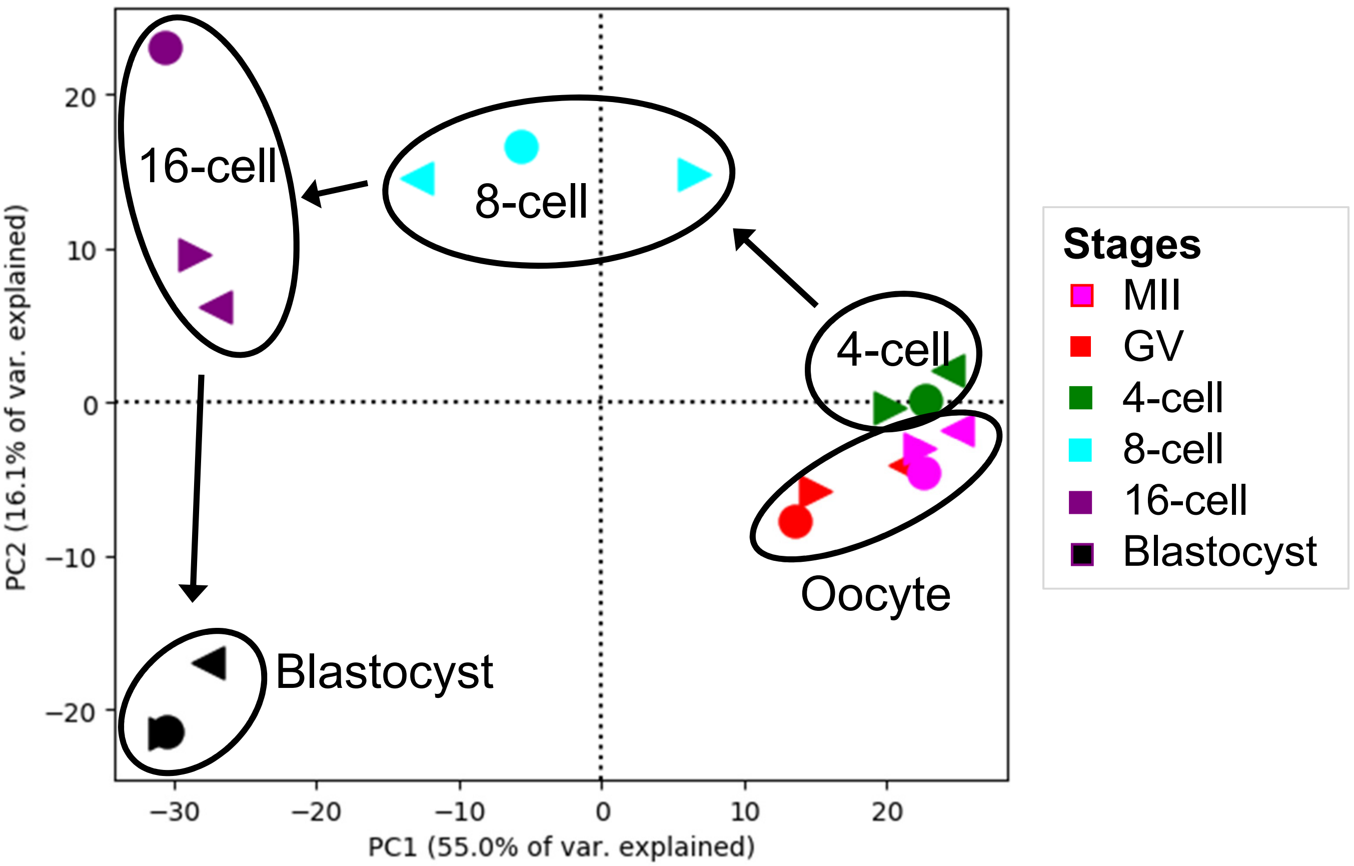


Figure S1. Principal components analysis (PCA) of RNA-seq read depth normalized by reads per kilobase million (RPKM) in 50 bp windows covering the whole genome. Sequencing data from Graf *et al,* (2014).

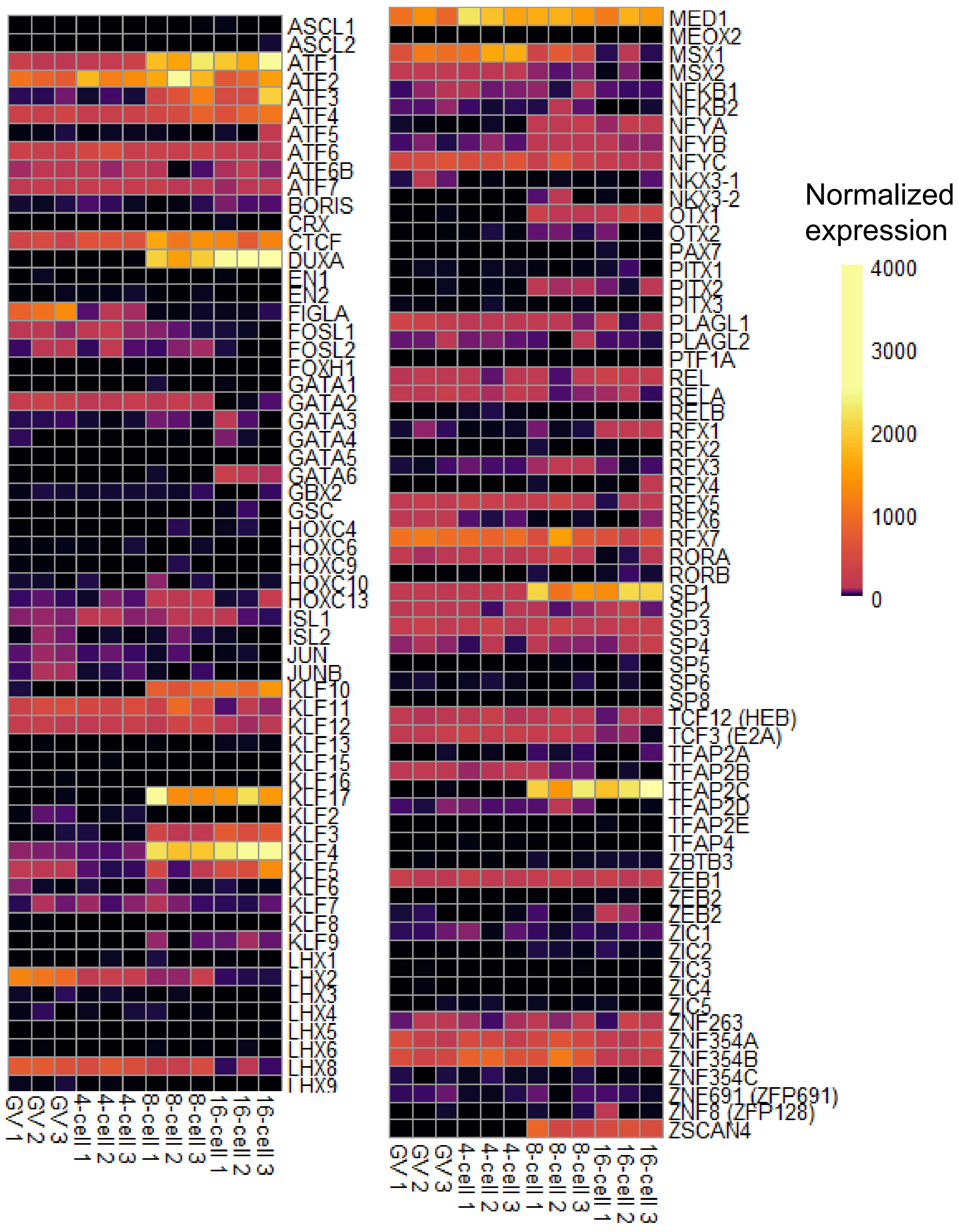


Figure S2. DESeq2 normalized expression of transcription factors with enriched binding motifs in distal open chromatin at some point during bovine preimplantation development. RNA-seq data from Graf *et al,* (2014).

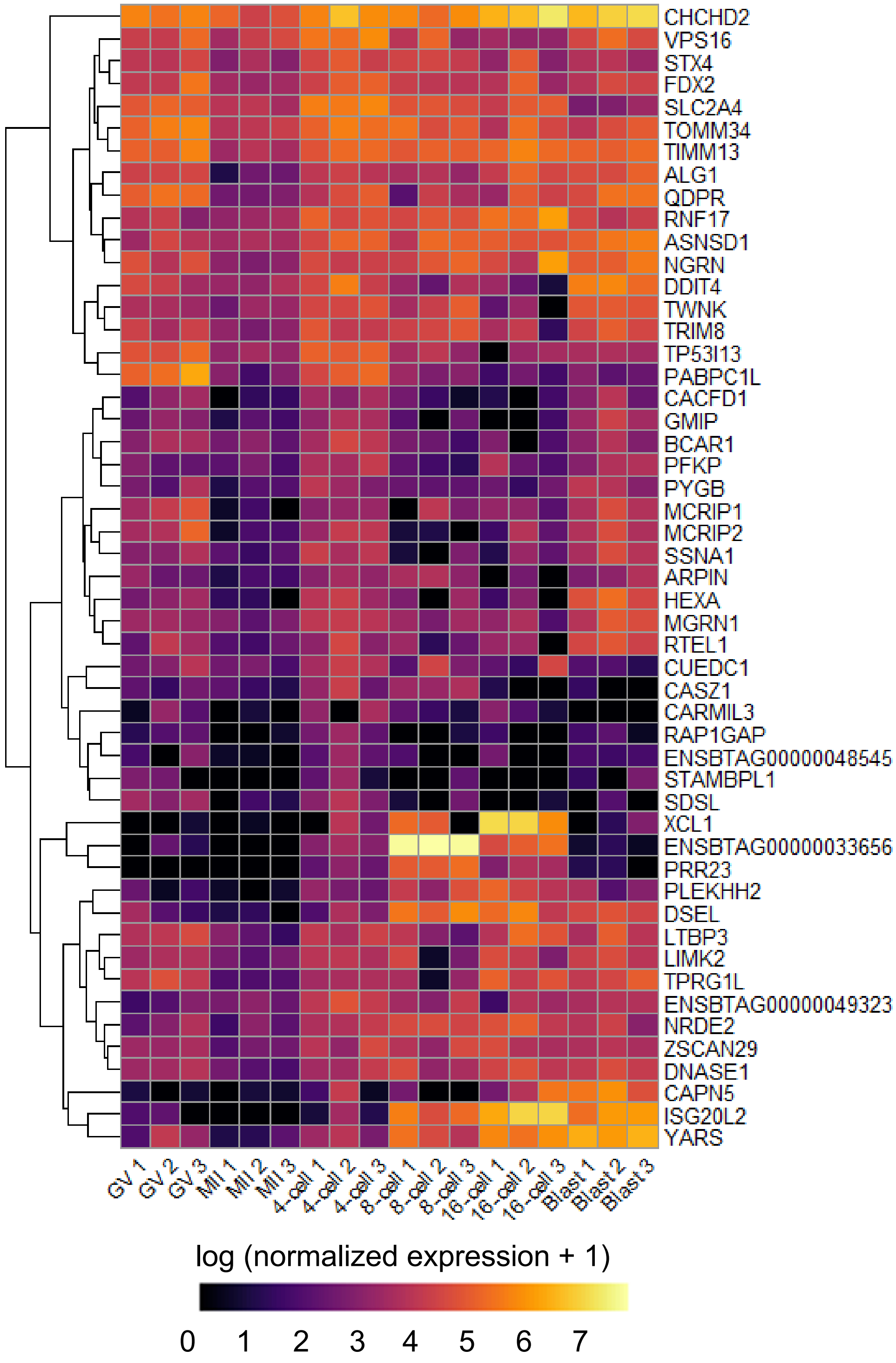


Figure S3. Minor EGA products. Log-transformed DESeq2 normalized expression of genes that were differentially upregulated in 4-cell embryos relative to MII oocytes (n=62 genes; adjusted $p < 0.05$; logFC > 2). RNA-seq data from Graf *et al,* (2014).

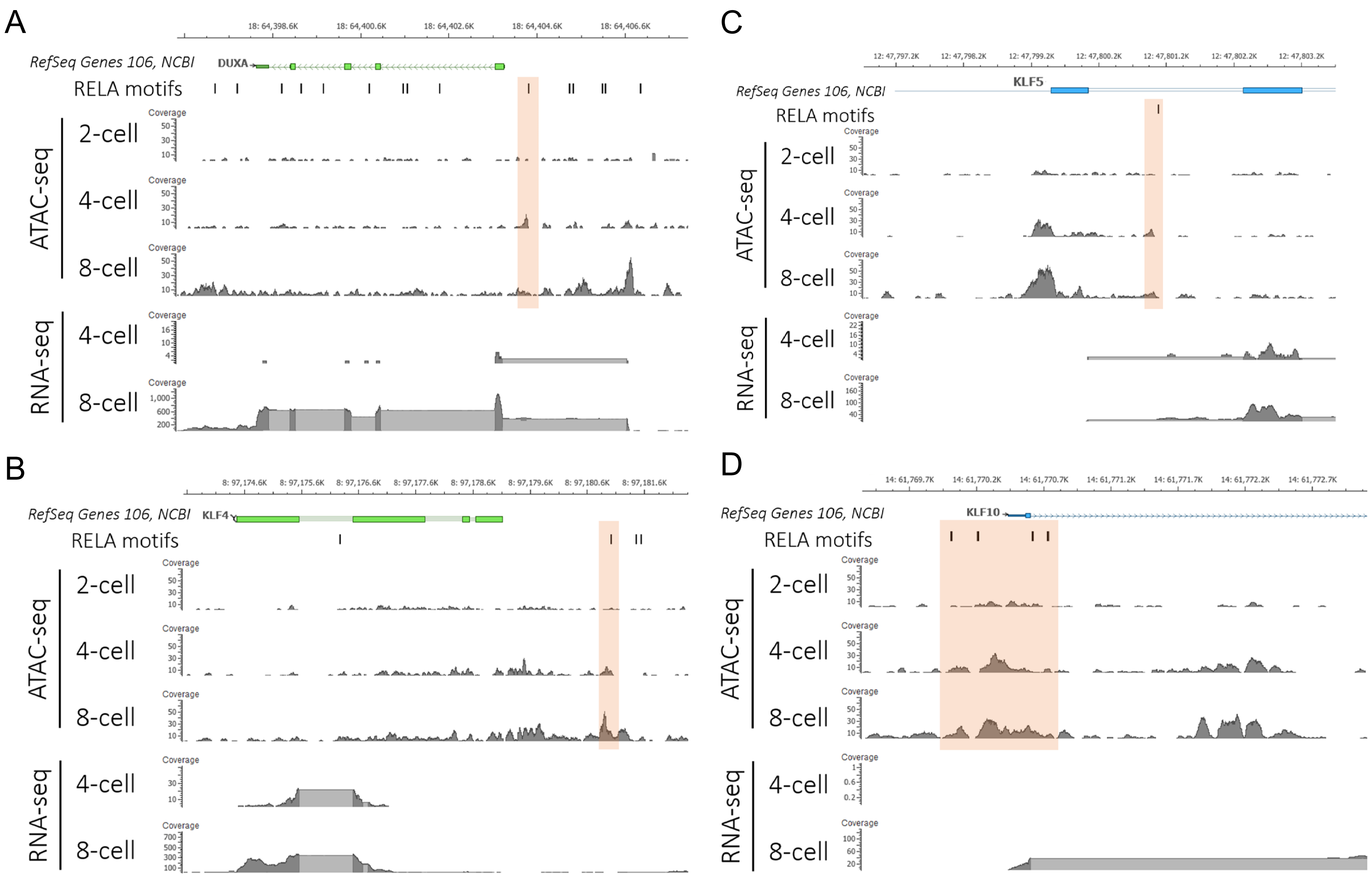


Figure S4. Accessible regions at key regulators of early development harbor RELA binding motifs. Read coverage is reported for each library and scaled to equal depth for all ATAC-seq libraries, each consisting of 30 million reads in total. Open chromatin in 4-cell embryos that overlapped RELA binding motifs highlighted in orange. A) DUXA; B) KLF5; C) KLF4; D) KLF10. RNA-seq data from Graf *et al,* (2014).

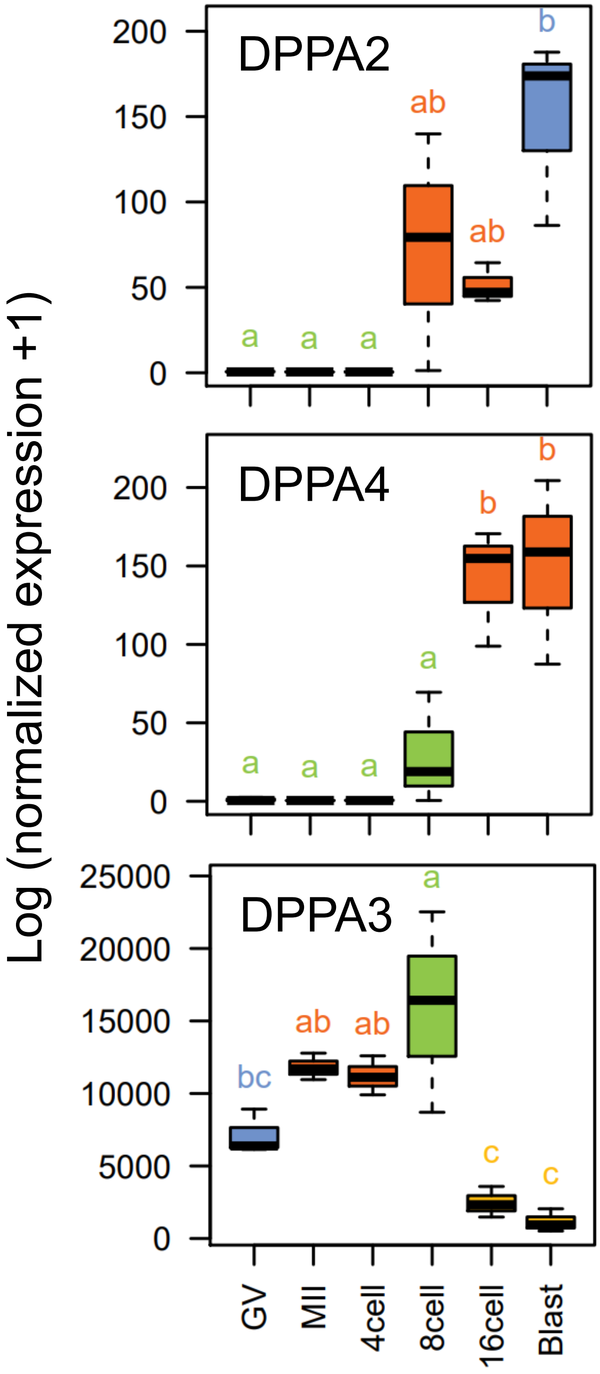


Figure S5. Expression profiles of DPPA genes during development. Expression values shown are log-transformed DESeq2-normalized counts. Colors and letters indicate statistical differences between stages, as calculated by one-way ANOVA with post-hoc Tukey test with a family-wise confidence level of 0.95. RNA-seq data from Graf *et al* (2014).

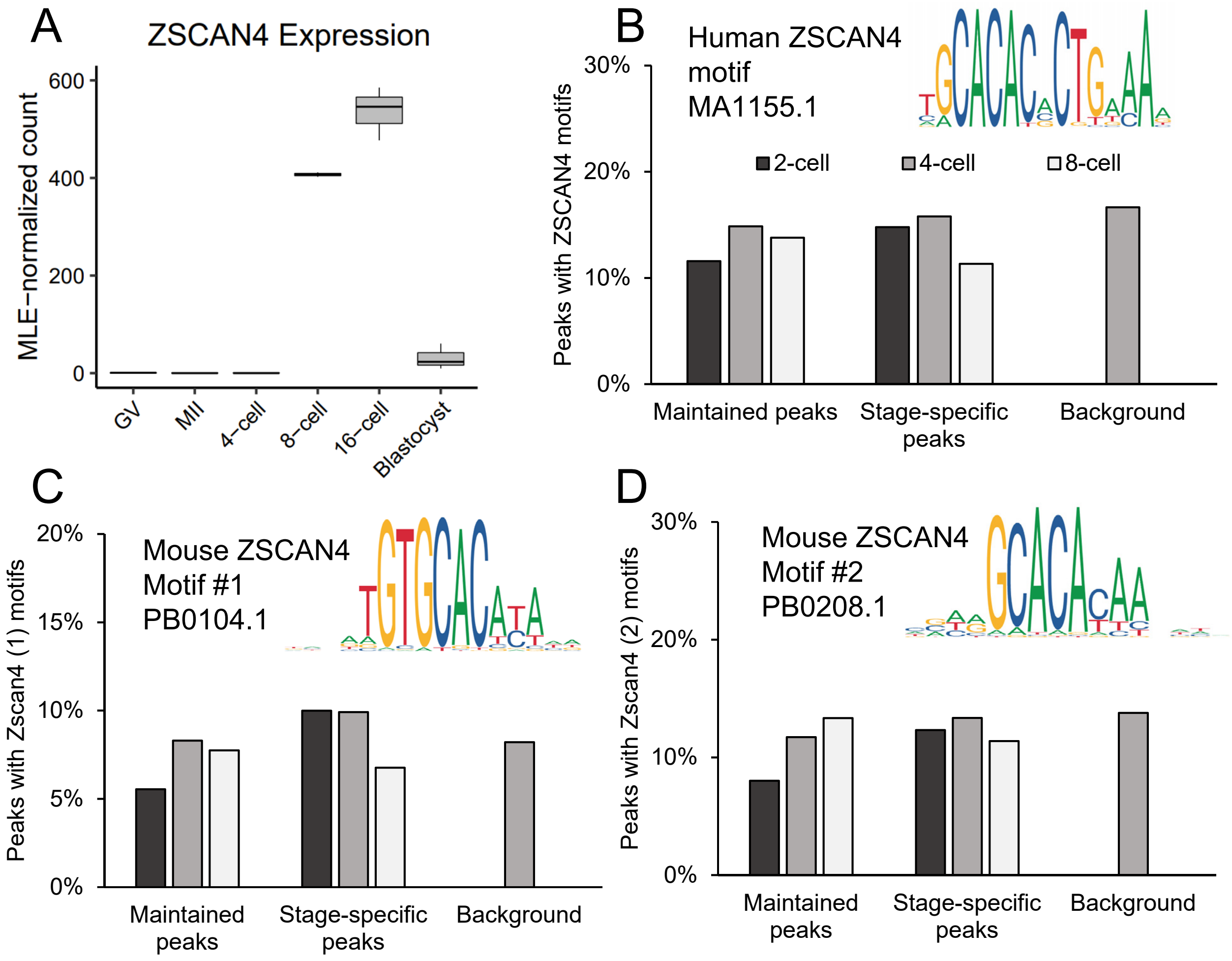


Figure S6. A) Enrichment of the ZSCAN4 motif in stage-specific and maintained peaks at the 2-, 4-, and 8-cell stages, relative to random background. B) DESeq2 normalized expression of ZSCAN4. RNA-seq data from Graf *et al* (2014).

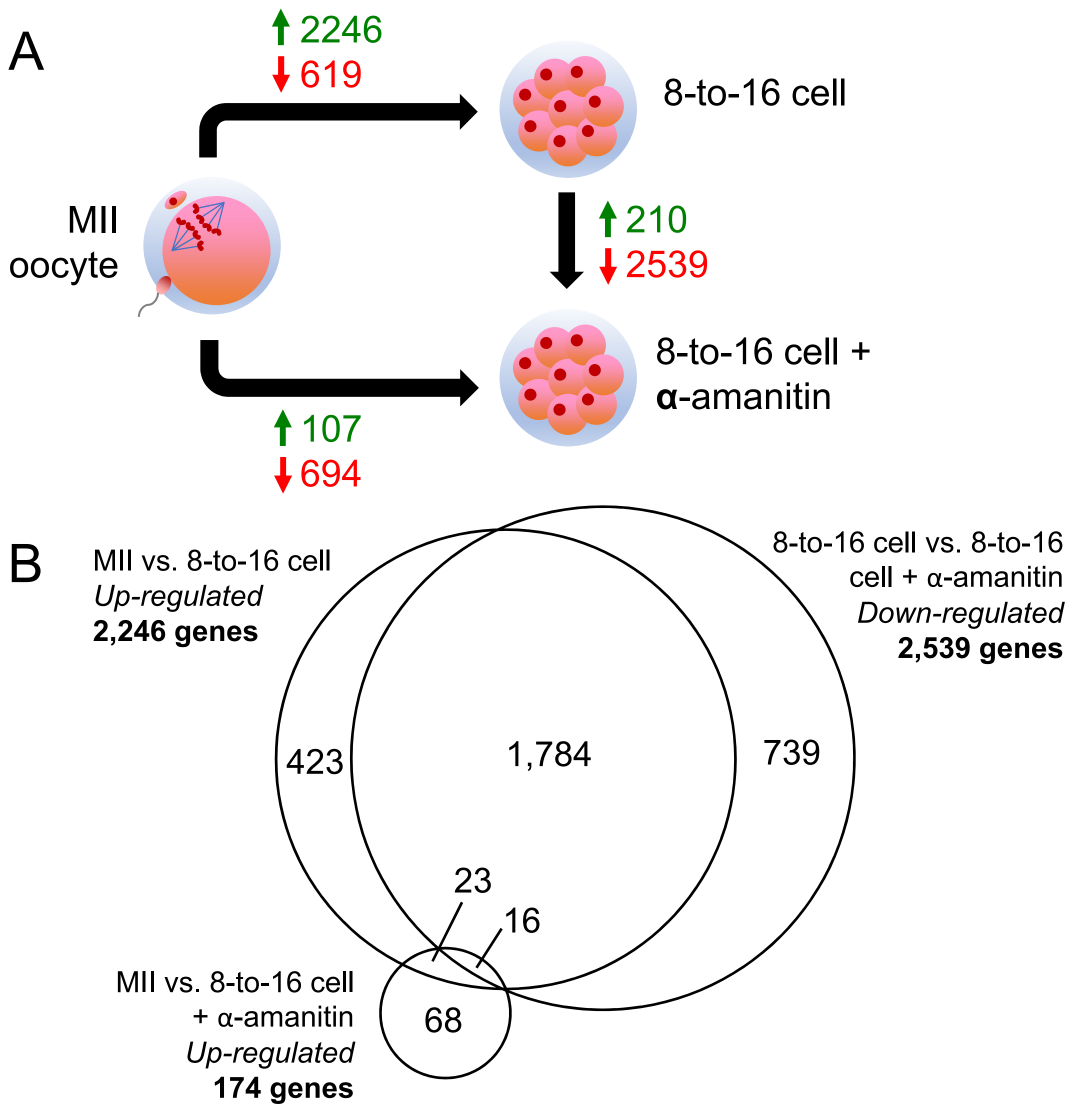


Figure S7. Analysis of RNA-seq data from Bogliotti *et al* (2019). A) Differentially expressed genes across developmental stages, which demonstrated an FDR < 0.01 and log2FC > 2. B) Venn diagram comparing DEG between different stages, allowing for the identification of 68 transcripts that were exclusively maternal in origin, and 1,784 transcripts that were exclusively embryonic in origin; i.e. EGA-specific genes.

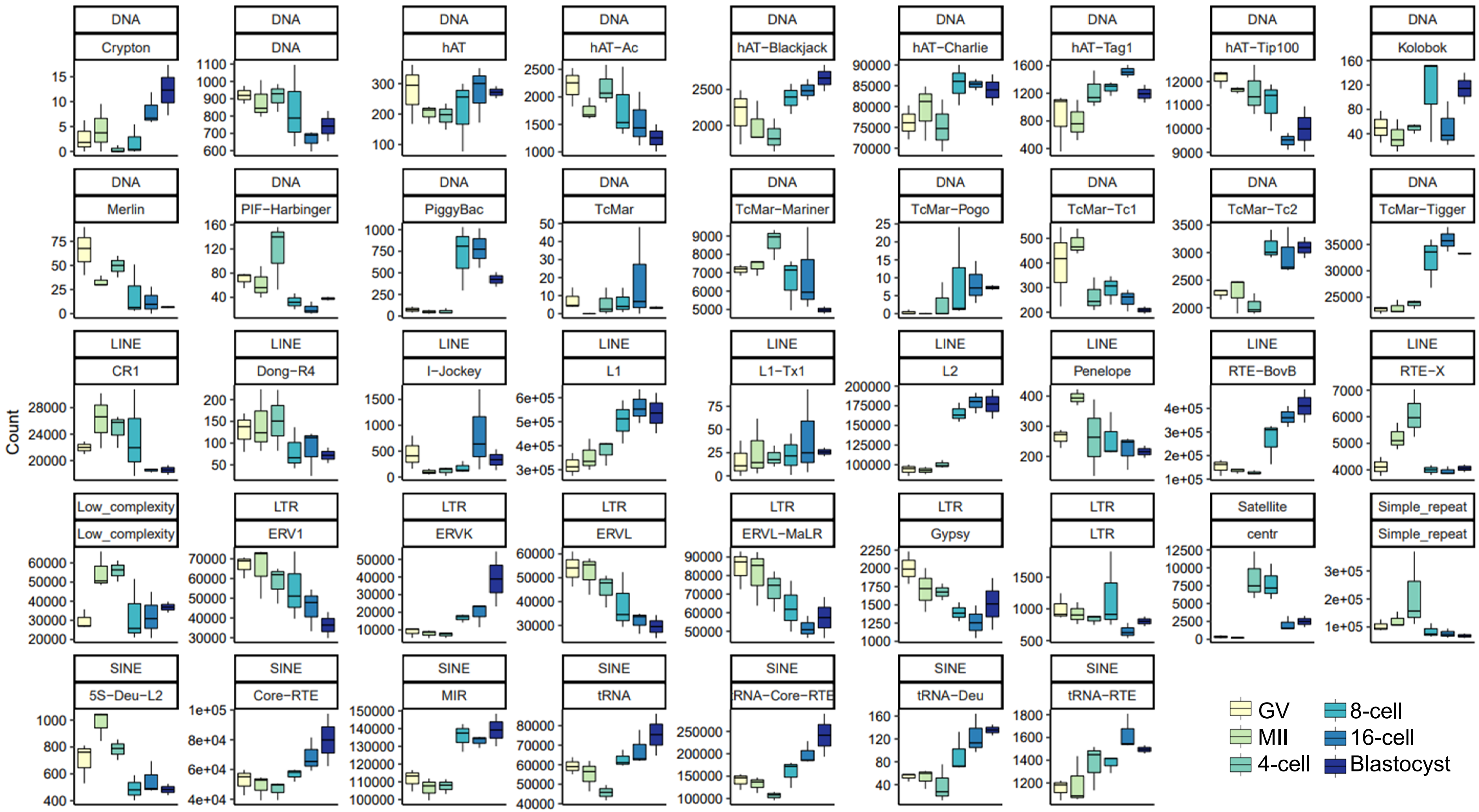


Figure S8. Normalized expression profiles of all LTR, LINE, and SINE repeat families throughout bovine preimplantation development. Transcriptomic data generated for bovine oocytes and embryos by Graf *et al* (2014).

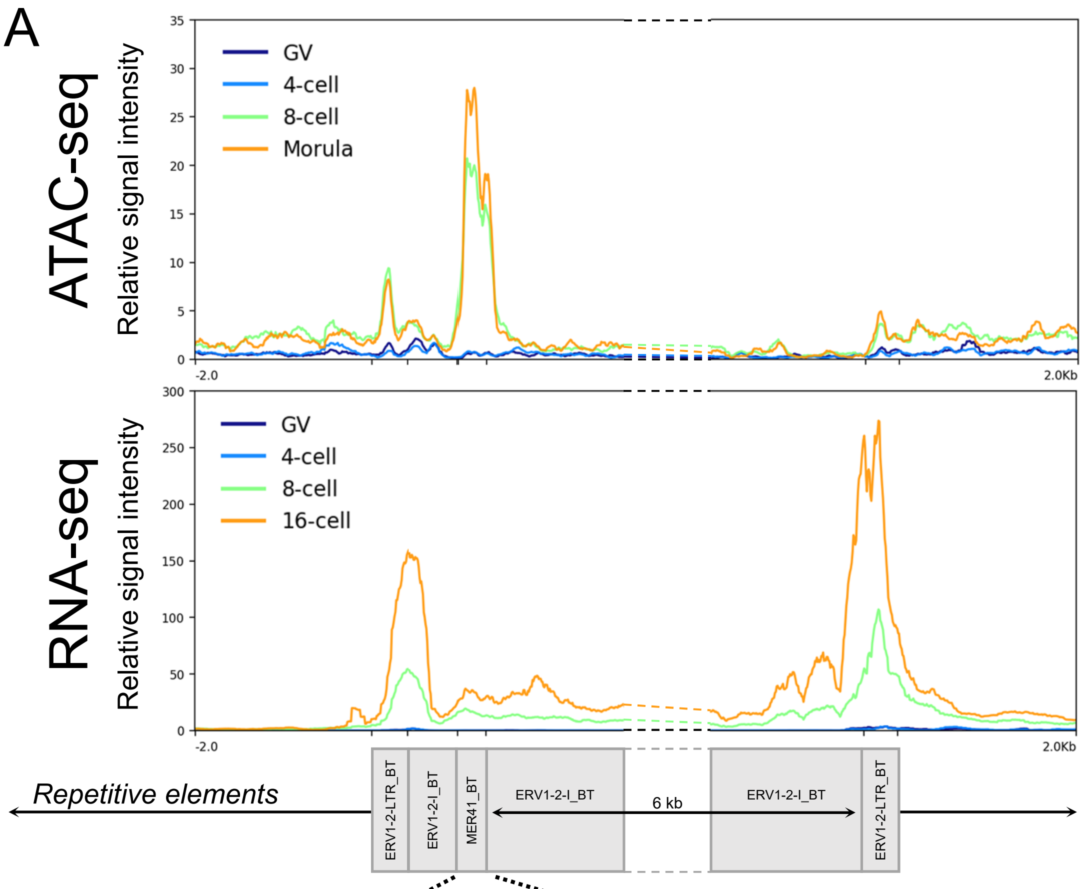


B

| Motif | P-value | Open chromatin in MER41 with motif (%) |
|---|---|---|
| POU5F1-SOX2-TCF-NANOG | 1e-20 | 29.4 |
| POU5F1 | 1e-13 | 33.3 |
| LHX2 | 1e-09 | 39.2 |
| NFY | 1e-07 | 33.3 |
| KLF4 | 1e-07 | 23.5 |
| OTX2 | 1e-07 | 31.4 |
| TEAD | 1e-06 | 31.4 |

Figure S9. A) Average relative ATAC-seq and RNA-seq signal intensity at 57 ERV1-2-I_BT elements flanked by ERV1-2-LTR_BT with an internal MER41_BT element. B) Top known motifs enriched in open chromatin at MER41_BT elements that accompany ERV1-2-I_BT and ERV1-2-LTR_BT elements. RNA-seq data from Graf *et al* (2014).

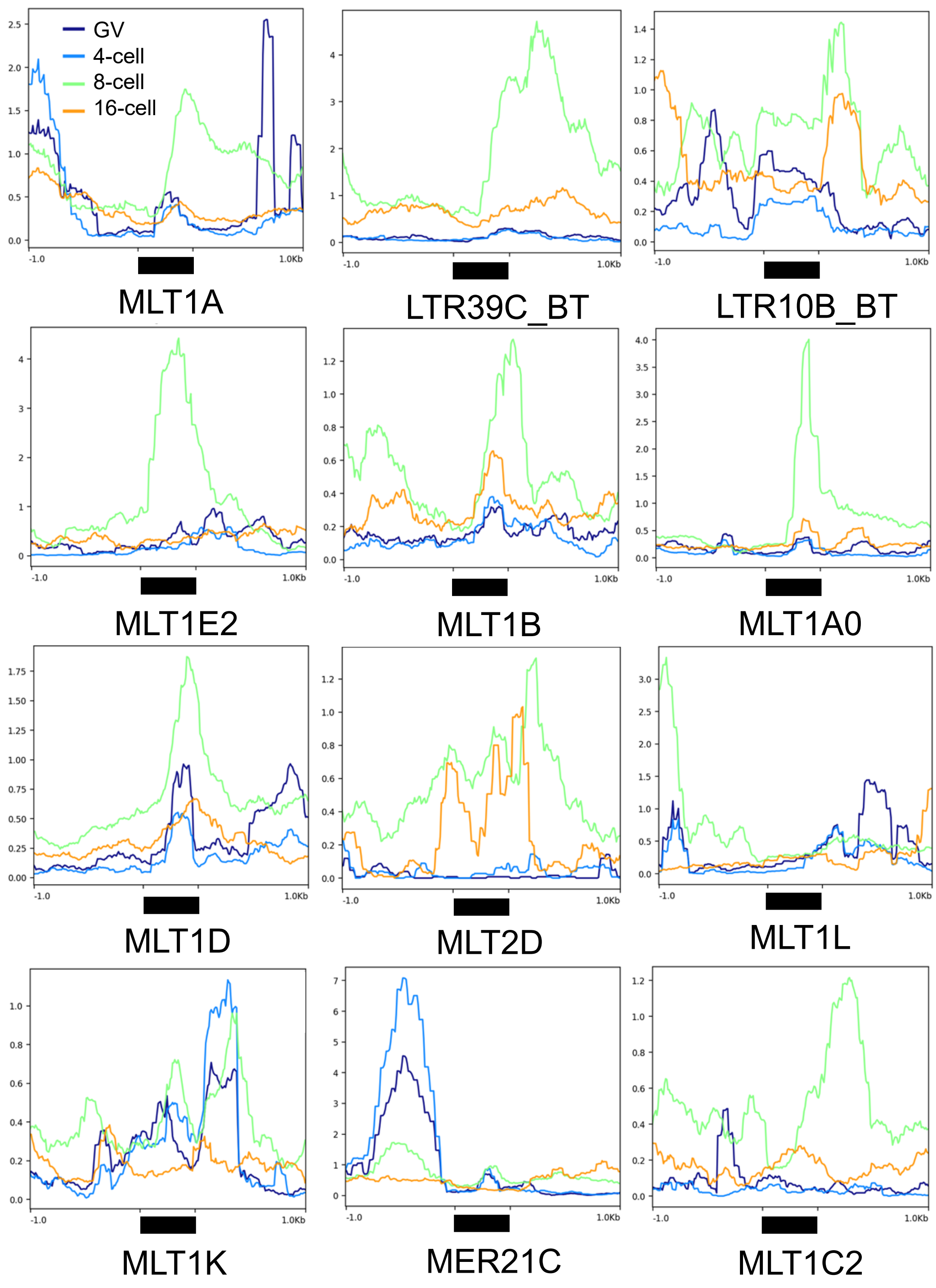


Figure S10. Normalized RNA-seq signal (RPKM) at repeats overlapping distal 8-cell ATAC-seq peaks harboring DUXA motifs. RNA-seq from Graf *et al* (2014).

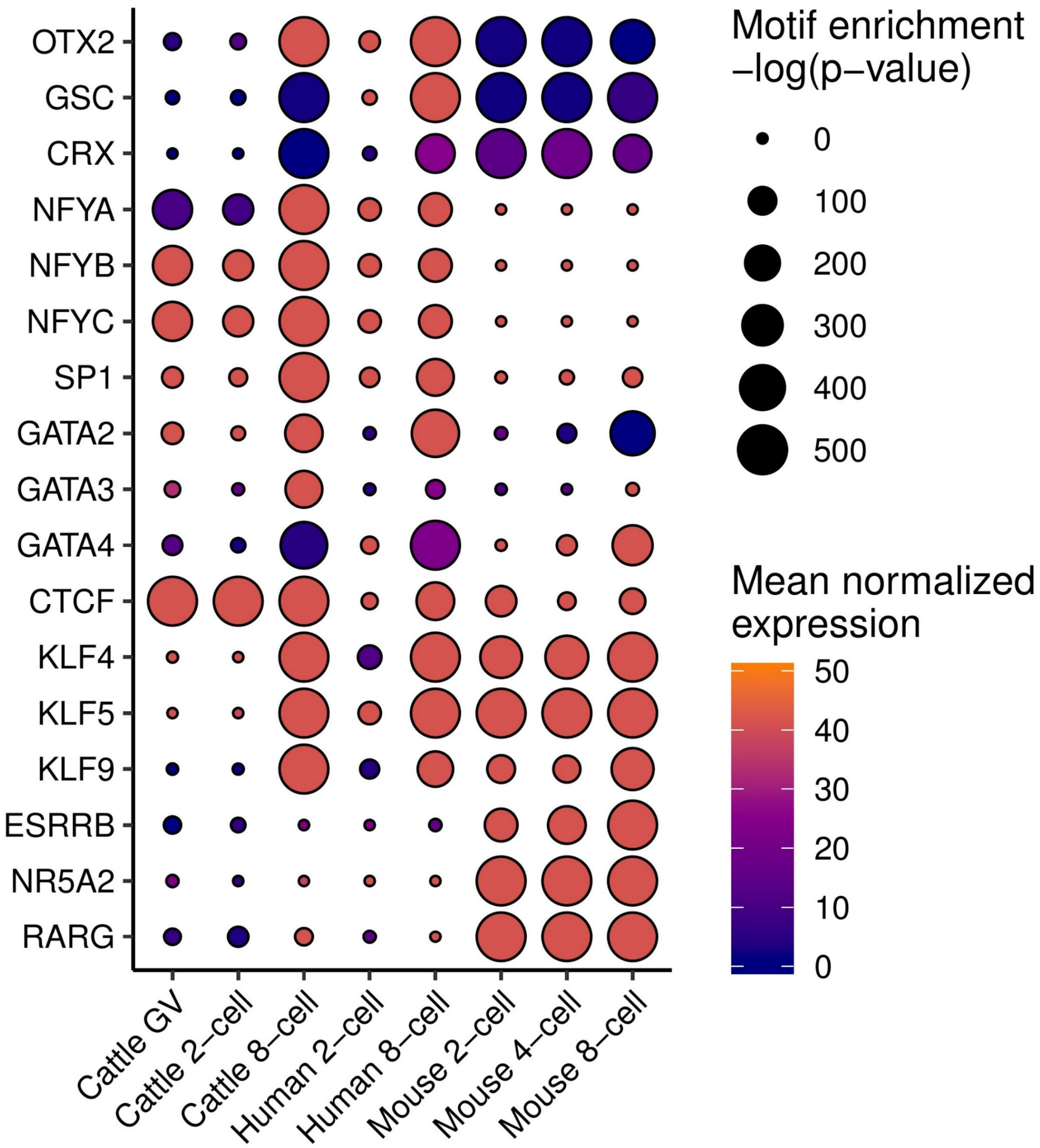


Figure S11. Inference of key regulatory factors during preimplantation development in cattle, human, and mouse, based on enrichment for TF binding factors in open chromatin and expression of the corresponding TF. Bovine RNA-seq data from Graf *et al* (2014); human ATAC-seq and RNA-seq data from Wu *et al* (2018); mouse ATAC-seq and RNA-seq from Wu *et al* (2016).

**Supplementary Tables**

Table S1. Sequencing depth of individual ATAC-seq libraries. Total raw sequencing reads, percent alignment to the genome (excluding the mitochondrial DNA), percent duplication among aligned reads, and total informative (non-duplicate, non-mitochondrial uniquely mapping) reads in each replicate that were available for peak calling.

| *Stage* | *Treatment* | *Rep.* | *Raw reads* | *Aligned reads (%)* | *Duplicate reads (%)* | *Informative reads* | *Total informative reads* |
|---|---|---|---|---|---|---|---|
| GV oocyte | Control | 1 | 237,846,070 | 56.92 | 90.71 | 7,520,394 | 30,191,548 |
| | | 2 | 156,780,590 | 76.35 | 89.22 | 7,597,123 | |
| | | 3 | 195,336,400 | 68.61 | 83.59 | 15,074,031 | |
| 2-cell | TBE | 1 | 462,735,044 | 43.21 | 95.96 | 3,649,130 | 19,536,740 |
| | | 2 | 67,607,200 | 59.97 | 77.60 | 5,792,638 | |
| | | 3 | 113,594,712 | 67.41 | 80.08 | 10,094,972 | |
| 2-cell | Control | 1 | 325,738,800 | 41.47 | 87.54 | 6,944,410 | 31,690,021 |
| | | 2 | 284,762,902 | 47.72 | 93.78 | 4,044,907 | |
| | | 3 | 107,281,710 | 72.97 | 61.13 | 20,700,704 | |
| 4-cell | TBE | 1 | 758,041,924 | 43.26 | 94.17 | 10,208,325 | 19,198,081 |
| | | 2 | 310,266,952 | 54.45 | 94.48 | 4,266,797 | |
| | | 3 | 51,133,698 | 49.88 | 71.15 | 4,722,959 | |
| 4-cell | Control | 1 | 744,499,540 | 50.11 | 92.99 | 14,825,045 | 36,653,439 |
| | | 2 | 178,231,726 | 57.57 | 76.83 | 14,011,042 | |
| | | 3 | 280,964,432 | 55.53 | 90.83 | 7,817,352 | |
| 8-cell | TBE | 1 | 134,460,968 | 91.69 | 58.74 | 36,786,855 | 163,235,550 |
| | | 2 | 538,922,446 | 85.69 | 65.42 | 116,170,135 | |
| | | 3 | 100,844,568 | 50.38 | 71.33 | 10,278,560 | |
| 8-cell | Control | 1 | 90,270,884 | 91.75 | 40.46 | 38,568,595 | 87,094,541 |
| | | 2 | 180,127,326 | 56.53 | 69.97 | 23,632,106 | |
| | | 3 | 216,383,872 | 52.26 | 71.43 | 24,893,840 | |
| Morula | Control | 1 | 127,179,516 | 73.94 | 81.43 | 13,744,880 | 64,586,045 |
| | | 2 | 160,860,156 | 66.12 | 67.21 | 28,915,042 | |
| | | 3 | 64,511,504 | 69.71 | 42.24 | 21,926,123 | |

Table S2. Spearman correlation coefficients calculated for replicate ATAC-seq libraries from genome-wide coverage, normalized for sequencing depth.

| | | *Spearman Correlation Coefficient* | | | |
|---|---|---|---|---|---|
| *Stage* | *Treatment* | *Rep. 1 – Rep. 2* | *Rep. 1 – Rep. 3* | *Rep. 2 – Rep. 3* | *Average* |
| GV | Control | 0.82 | 0.84 | 0.84 | 0.83 |
| 2-cell | Control | 0.63 | 0.70 | 0.69 | 0.67 |
| 4-cell | Control | 0.80 | 0.77 | 0.75 | 0.77 |
| 8-cell | Control | 0.89 | 0.88 | 0.86 | 0.88 |
| Morula | Control | 0.87 | 0.86 | 0.91 | 0.88 |
| 2-cell | TBE | 0.54 | 0.53 | 0.74 | 0.60 |
| 4-cell | TBE | 0.63 | 0.70 | 0.59 | 0.64 |
| 8-cell | TBE | 0.90 | 0.86 | 0.83 | 0.86 |

Table S3. Top five *de novo* enriched motifs found in intergenic open chromatin that was either lost in 2-cell embryos or gained in 4-cell, 8-cell, or morula-stage embryos. Known motifs with a match score > 0.6 reported for each *de novo* motif.

| | *Motif* | *Matching known motifs* | *P-value* | *Loci with motif (%)* | *Background (%)* |
|---|---|---|---|---|---|
| Closed from GV to 2-cell | 1 | BORIS, CTCF, SP4, ZIC1 | 1e-656 | 3.56 | 0.75 |
| | 2 | MEOX2, EN2, EN1, LHX9, MSX1, LHX8, MEOX1, GBX2 | 1e-235 | 43.64 | 36.78 |
| | 3 | PTF1A, HEB, AP4, E2A, ASCL1, ZEB1, TCF12, FIGLA, MYOG | 1e-230 | 28.47 | 22.48 |
| | 4 | FOSL1, JUNB, BATF, FOSL2, AP1, ATF3, JUN-AP1 | 1e-226 | 5.10 | 2.61 |
| | 5 | RFX1, RFX5, X-box, RFX4, RFX2, RFX3, RFX | 1e-216 | 8.83 | 5.50 |
| Opened from 2- to 4-cell | 1 | BORIS, CTCF, SP4, ZIC1, ZIC4 | 1e-830 | 5.01 | 0.80 |
| | 2 | NFkB-p65, NFKB2, REL, NFkB-p50,p52, RELA, NFKB1 | 1e-132 | 10.49 | 7.04 |
| | 3 | ZFP128, PLAGL1, ZNF354C, ZNF263, CRX, FOXH1 | 1e-126 | 0.22 | 0.00 |
| | 4 | NKX3-1, ISL2, NKX3-2 | 1e-113 | 0.24 | 0.01 |
| | 5 | ZFP691, ZNF263, MED1, SOX8, ZBTB3 | 1e-109 | 0.95 | 0.22 |
| Opened from 4- to 8-cell | 1 | DUX4, CRX, DMBX1, PITX1, PITX3, PAX7, GSC | 1e-5781 | 35.17 | 9.81 |
| | 2 | KLF5, KLF4, KLF1, GC-box, KLF14, KLF7, SP3 | 1e-1687 | 14.52 | 4.81 |
| | 3 | NFYB, NFY, CCAAT-box, NFYA, DUX | 1e-1543 | 11.57 | 3.44 |
| | 4 | EWS-ERG, HOXC9, POU6F2 | 1e-1253 | 2.43 | 0.12 |
| | 5 | RORgt, RORA | 1e-999 | 1.90 | 0.09 |
| Opened from 8-cell to morula | 1 | KLF4, KLF5, KLF1, KLF7, GC-box, SP3, KLF14 | 1e-1375 | 20.21 | 9.84 |
| | 2 | GATA4, GATA3, GATA5, GATA2, GATA1, GATA6 | 1e-1194 | 30.59 | 18.60 |
| | 3 | CRX, OTX2, PITX3, PITX1, GSC, DMBX1, PITX2, OBOX6 | 1e-878 | 15.11 | 7.72 |
| | 4 | AP-2gamma, AP-2alpha, TFAP2E, TFAP2C, TFAP2A, TFAP2B | 1e-864 | 15.85 | 8.32 |
| | 5 | BORIS, CTCF, SP4, ZIC1, ZIC4 | 1e-736 | 4.03 | 1.03 |

Table S4. Top enriched known motifs in loci that opened at the A) 2-cell, B) 4-cell, or C) 8-cell stages and which remained accessible up the morula stage.

| | | *Motif* | *P-value* | *Peaks with motif (%)* |
|---|---|---|---|---|
| A | Maintained 2-cell open chromatin | NFY | 1e-306 | 31.47 |
| | | CTCF | 1e-158 | 13.54 |
| | | SP1 | 1e-152 | 41.11 |
| | | GFX | 1e-126 | 4.15 |
| | | ETS | 1e-109 | 21.62 |
| | | ELK1 | 1e-98 | 34.47 |
| | | ZBTB33 | 1e-98 | 8.28 |
| B | Maintained 4-cell open chromatin | CTCF | 1e-2586 | 27.26 |
| | | BORIS | 1e-1571 | 34.77 |
| | | SP1 | 1e-229 | 29.07 |
| | | NFY | 1e-215 | 23.77 |
| | | ETS | 1e-167 | 16.72 |
| | | KLF9 | 1e-156 | 27.69 |
| | | NRF1 | 1e-147 | 12.95 |
| C | Maintained 8-cell open chromatin | OTX2 | 1e-4190 | 45.31 |
| | | GSC | 1e-3882 | 55.26 |
| | | CTCF | 1e-2287 | 10.24 |
| | | KLF5 | 1e-1782 | 37.45 |
| | | CRX | 1e-1758 | 71.94 |
| | | PHOX2A | 1e-1734 | 24.04 |
| | | KLF4 | 1e-1595 | 15.48 |

Table S5. Retrotransposon enrichment in 8-cell intergenic open chromatin harboring DUXA motifs (n=16,804 loci). Retrotransposons that appeared in at least 100 open chromatin regions reported.

| *Repeat* | *Distal 8-cell peaks with DUXA motif* | *Background* | *Log ratio* |
|---|---|---|---|
| MLT1A | 507 | 32 | 2.763 |
| LTR39C_BT | 125 | 14 | 2.189 |
| LTR10B_BT | 239 | 28 | 2.144 |
| MLT1E2 | 101 | 16 | 1.843 |
| MLT1B | 404 | 67 | 1.797 |
| MLT1A0 | 443 | 74 | 1.790 |
| MLT1D | 512 | 87 | 1.772 |
| MLT2D | 108 | 19 | 1.738 |
| MLT1L | 185 | 34 | 1.694 |
| MLT1K | 311 | 58 | 1.679 |
| MER21C | 150 | 45 | 1.204 |
| MLT1C2 | 197 | 65 | 1.109 |
